# Performance Evaluation of Ciprofloxacin loaded Micellar Bioconjugate Carrier for Eradication of Bacterial Biofilms

**DOI:** 10.1101/2024.12.25.630303

**Authors:** Chandrika Gupta, Sudip Kumar Ghosh, Ramkrishna Sen

## Abstract

This study reports the utilization of a Curcumin-Exopolysaccharide (Cur-EPS) micellar bioconjugate comprising acid-labile ester bond as a carrier for ciprofloxacin (Cip) to improve its ability to eradicate mature bacterial biofilms. The most significant attributes of Cip-loaded Cur-EPS micelles, including hydrodynamic particle size, polydispersity index (PDI), and encapsulation efficiency (EE), were statistically optimized to be 305.43 ± 1.47 nm, 0.38 ± 1.23, and 84.50 ± 2.01 %, respectively. At an acidic pH of 5.6, the pH-responsive release of Cip from Cur-EPS micelles was 82.51 ± 2.95 % over a duration of 100 h, but it was reduced to 39.10 ± 1.55 % at a pH of 7.4 over the same period. This pH-dependent Cip release may be attributed to the destabilisation of Cur-EPS micelles due to hydrolysis of the succinic acid linker in the acidic pH range. Thus, the acidic pH microenvironment of biofilms formed by pathogenic bacterial strains, such as methicillin resistant *Staphylococcus aureus* (MRSA) and *Pseudomonas aeruginosa*, enhances the pH-responsive release of Cip from Cur-EPS micelles, thereby leading to the eradication of mature biofilms. This enhanced antibiofilm effect of Cip-Cur-EPS, in contrast to the free form of Cip, was evident from the significant decrease in the biofilm biomass. Furthermore, mature biofilms that formed on the surface of the urinary catheter were almost eradicated when treated in vitro with Cip-Cur-EPS for 72 h in the presence of artificial urine media (AUM). Moreover, Cip-Cur-EPS could also eradicate the biofilms formed by *P. aeruginosa* and MRSA in a catheterized bladder model employing recirculated artificial urine. Overall, this study demonstrated the antibiofilm potential of Cip-Cur-EPS micelles, rendering them suitable therapeutic agents for addressing biofilm-associated infections in clinical settings.

**Graphical abstract:** 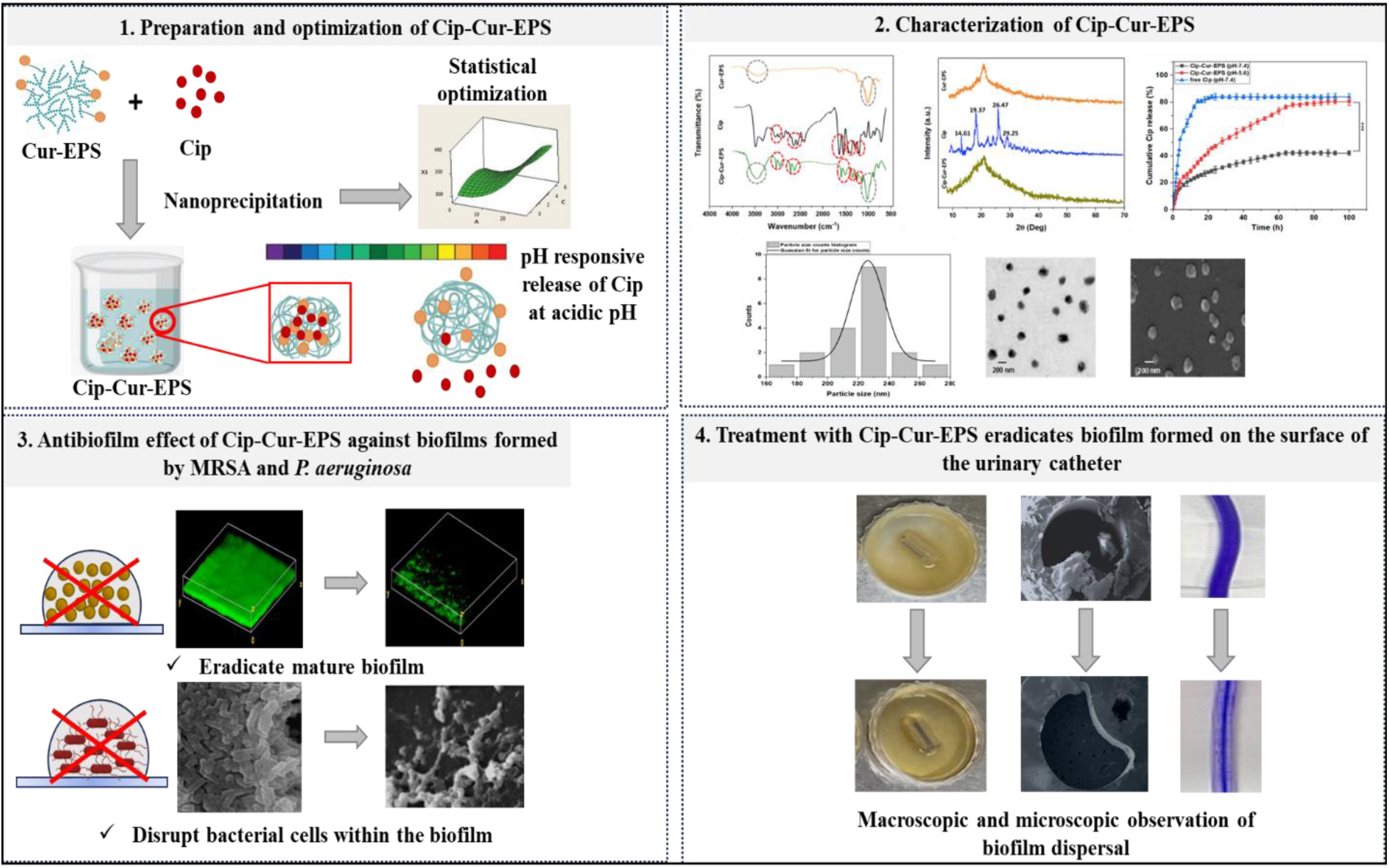

## 1. Introduction

Bacterial biofilm-associated infections pose major concerns globally in the healthcare sector. Biofilm infections commonly occur in association with the medical devices such as, prosthetic valves, implants, contact lenses, cerebrospinal fluid shunts and catheters. They can also be associated with tissue related infections such as skin infections, urinary tract infections cystic fibrosis and endocarditis, etc. (1)

As per the reports published by the National Institutes of Health (NIH), about 80 % of infectious diseases in patients are related to the biofilms. Therefore, formation of biofilms is recognized as the major cause for various persistent and chronic bacterial infections. (2) Over 2 million individuals are affected by biofilm related infections throughout the world. Further projections indicate a significant increase in occurrences of such infections without adequate intervention. (3)

Biofilms are microbial aggregates that are enveloped within the extracellular polymeric substances, which forms a matrix surrounding the cells. Extracellular polymeric substances primarily consist of water, proteins, polysaccharides, extracellular DNA, and lipids, which significantly contribute to the formation of biofilm matrix. Therefore, it plays a major role in enabling cellular interaction and adhesion on the abiotic and biotic surfaces. The assembly of bacterial cells in biofilm facilitates the establishment of a stable and mutually beneficial collaboration between cells, called quorum sending, leading to the development of a robust microbiome. (4) Numerous investigations have demonstrated that extracellular polymeric substances produced by the pathogenic bacterial strains creates a physical barrier, which impedes penetration of the antibiotics. (5) Therefore, bacterial cells embedded within the biofilms acquire adaptive resistance, which plays a crucial role in bacterial pathogenicity. (6) Moreover, mature biofilms further disassemble, leading to the release of individual cells and intact biofilm fragments (sloughing) into the bloodstream and surrounding tissues promoting recolonization of bacterial cells. Therefore, biofilm associated bacterial infections throw a significant challenge in eradicating biofilms due to their ability to shift from acute to chronic stages, leading to severe consequences. (7) Traditional therapy for treating the biofilm related infections typically requires treatment with high doses of antibiotics over a long period, thereby fostering antimicrobial resistance. Such treatments are insufficient to entirely eradicate the biofilms, leading to recurrent infections. Moreover, the elevated doses of antibiotics also lead to systemic drug toxicity. The most prevalent side effects that are reported during antibiotic treatment includes physical discomfort, hypersensitivity and unfavourable impact on the gastrointestinal, renal, neurological, pulmonary, cardiac, and hepatic systems. (8) Therefore, alternative treatment approach has been ventured to efficiently eradicate persistent biofilm infections without detrimental side effects.

The antimicrobial delivery systems in the form of carriers in sub-micron range from 0.1 – 0.5 µm, have been employed to treat infectious biofilms and can alleviate the antibiotic resistance and their unwanted side effects. (9) Such delivery systems can be further tailored depending on the distinct physiological feature specifically present in the biofilm microenvironment in order to enable efficient release of antibiotics within biofilms. The presence of the anoxic environment within the biofilm enables the production of the acidic byproducts via anaerobic fermentation, consequently lowering the pH within the biofilm (pH < 6) compared to the normal physiological pH (7.4). This internal stimulus can be further leveraged for the release of antimicrobials within the biofilms. (9) Therefore, antibiotic nanocarriers endowed with pH-responsive properties can facilitate targeted antimicrobial release within the biofilms. (10)

We have previously reported the synthesis and characterization of pH-responsive Cur-EPS micelles by including a succinic acid spacer, which offers an acid labile ester linkage. The Cur-EPS micelles were biocompatible with significant antibiofilm activity against *P. aeruginosa*, *E. coli*, *S. aureus*, *S. typhimurium,* and *S. marcescens*. (11) The Cur-EPS micelles can be utilized as a micellar bioconjugate carrier to encapsulate drugs to improve its bioavailability at the biofilm infection site. Therefore, in the present study, Cur-EPS micelles were utilized as the carrier to encapsulate Ciprofloxacin (Cip), which is a broad-spectrum antibiotic belonging to the second-generation fluroquinolone comprising of cyclopropyl, carboxylic acid, fluoro and piperazin-1-yl substituents at 1^st^, 3^rd^, 6^th^ and 7^th^ positions respectively. It is a topoisomerase IV inhibitor approved by Food and Drug Administration (FDA) respectively, for the treatment of several bacterial illnesses. (12)

The effective treatment relies on the solubility of Cip and its efficient distribution to the infection site. However, Cip has inadequate solubility at physiological pH of 7.4, resulting in low bioavailability. Therefore, biocompatible Cur-EPS micelles were employed as a pH-responsive carrier for Cip, which can enhance the solubility of Cip and also enable its release in order to effectively eradicate the biofilms. Furthermore, effectiveness of these micelles were determined to eradicate mature biofilms formed on the surface of urinary catheters. Several patients become vulnerable to infections due to the repeated insertion of indwelling catheters, since biofilms may develop on their surfaces influenced by various factors such as infection history of the patient, immune function, microbiota, and the specific pathogens existing in the hospital. (13)

The present study was aimed to prepare Cip loaded Cur-EPS micelles. The formulation parameters for preparation of Cip-loaded polymeric micelles were statistically optimized to achieve the optimal mean particle size, PDI, and EE in order to demonstrate significant antibiofilm effect. Moreover, the most optimal formulation was characterized in terms of its structure and morphology, *in vitro* release of Cip and storage stability. Furthermore, the *in vitro* antibiofilm activities of Cip-Cur-EPS were examined using the biofilms formed by Gram-negative and Gram-positive bacteria. The antibiofilm effect of Cip loaded Cur-EPS micelles were evaluated in terms of reduction in biofilm biomass. Following the demonstration of the antibiofilm properties of Cip-loaded Cur-EPS micelles, their practical applicability was subsequently assessed to eradicate mature biofilms formed by *P. aeruginosa* and MRSA on the surface of urinary catheter in AUM.

## 2. Materials and methods

### 2.1. Materials

The Cur-EPS conjugate was synthesized and characterized in our laboratory and was reported earlier. (11) Ciprofloxacin (Cip, Mw = 331.34, purity ≥ 98%), Tween 80, Dimethyl sulfoxide (DMSO), crystal violet and acridine orange were purchased form Merck, Bangalore, India. 2,3,5-triphenyl tetrazolium chloride (TTC) was procured from LOBA Chemie, Mumbai, India. Tryptic soy broth (TSB), Muller Hinton Broth (MHB) and agar powder were procured from (Himedia, Mumbai, India).

### 2.2. Preparation of Cip-Cur-EPS

Cip-loaded Cur-EPS micelles were prepared by employing nanoprecipitation process, adhering to the previously reported method with minor modification. (14) Precise quantities of both Cur-EPS conjugate and Cip were dissolved in a solvent that was miscible with water, such as DMSO. The organic solution of Cur-EPS and Cip were added dropwise (0.5 mL/min) into aqueous solution containing a non-ionic surfactant (Tween-80) under stirring condition at 300 rpm for 12 h. Subsequently, the suspension was dialysis against deionized water using a dialysis membrane with a molecular weight cut-off (MWCO) of 3500 Da for 24 h followed by centrifugation at 10,000 rpm to remove non-encapsulated Cip and remaining DMSO. The DMSO was further removed by centrifuge concentrator (Eppendorf Vacufuge plus, USA). The micelle was resuspended in Milli Q water was subjected to an additional filtration process using a sterile syringe filter (0.22 μm pore size, Millex®, Merck, Mumbai, India) to eliminate remaining free Cip. The resulting Cip loaded micelle solution was then freeze dried for the purpose of preservation and characterization. The blank nanoparticles were also synthesized without loading Cip by following similar procedure.

### 2.3. Optimization of Cip-Cur-EPS micelles by experimental design

The Cip loaded Cur-EPS nanoparticles were optimized by implementing the design of experiments (DoE) using Minitab® 22.1 software (Minitab, LLC Pennsylvania, USA). The optimization process entailed design formulations to reduce number of trials and analyse response surfaces to comprehend the effects of the factors on desired response. (15) From the preliminary experiments it was evident that the variables, such as Cur-EPS concentration, surfactant concentration, and aqueous phase to organic phase ratio had influence on the hydrodynamic particle size, PDI and EE of the Cip-Cur-EPS micelles. Therefore, 3-factor and 3-response central composite design–response surface methodology (CCD–RSM) were employed for the optimization of Cip loaded micellar nanoparticles employing formulation variables such as Cur-EPS concentration (A), Tween 80 (B), and water:DMSO (C), as independent variables (factors). Further their effects were evaluated on particle size, PDI (polydispersity index) and EE (encapsulation efficiency) as dependent variables (responses). In order to minimize the number of experiments, a full factorial central design with three independent variables and five levels (two axial points, two cube points and a central point) were implemented. The information regarding the experimental design is indicated in Table 1. Additionally, the design incorporated a total of twenty experimental trials fitted into a multiple linear regression model. The regression coefficient (R²) was utilized for the identification of the model fit. Furthermore, to demonstrate the statistical validity of the generated equations, ANOVA was applied. The optimal experimental model, encompassing main effects, interactions, and quadratic terms, were selected based on a comparative analysis of various statistical parameters, such as the coefficient of variation, multiple correlation coefficient (R²), predicted R² (Pred.R²) and adjusted R² (Adj.R²). The significance level was set at p-value < 0.05. A mathematical correlation between independent factors and the responses were established by response surface regression analysis using Minitab software. The standard form of a second-order polynomial regression model with three variables were employed to demonstrate the response behaviour, as indicated in the following equation:

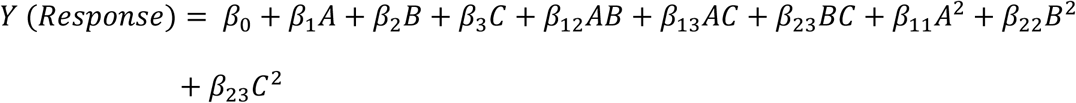

where *A*, *B*, and *C* are the coded value of independent variables; *β_0_* is the constant; *β_1_*, *β_2_*, and *β_3_* are linear coefficients; *β_12_*, *β_13_*, and *β_23_* are interaction coefficients between the independent variables; *β_11_*, *β_22_*, and *β_33_* are quadratic coefficients. Furthermore, *A*, *B*, and *C* represent the main effects, average value of changing factor one at a time, *AB* and *AC* and *BC* represent the interaction terms and *A^2^*, *B^2^* and *C^2^*are the polynomial terms.

**Table 1.**
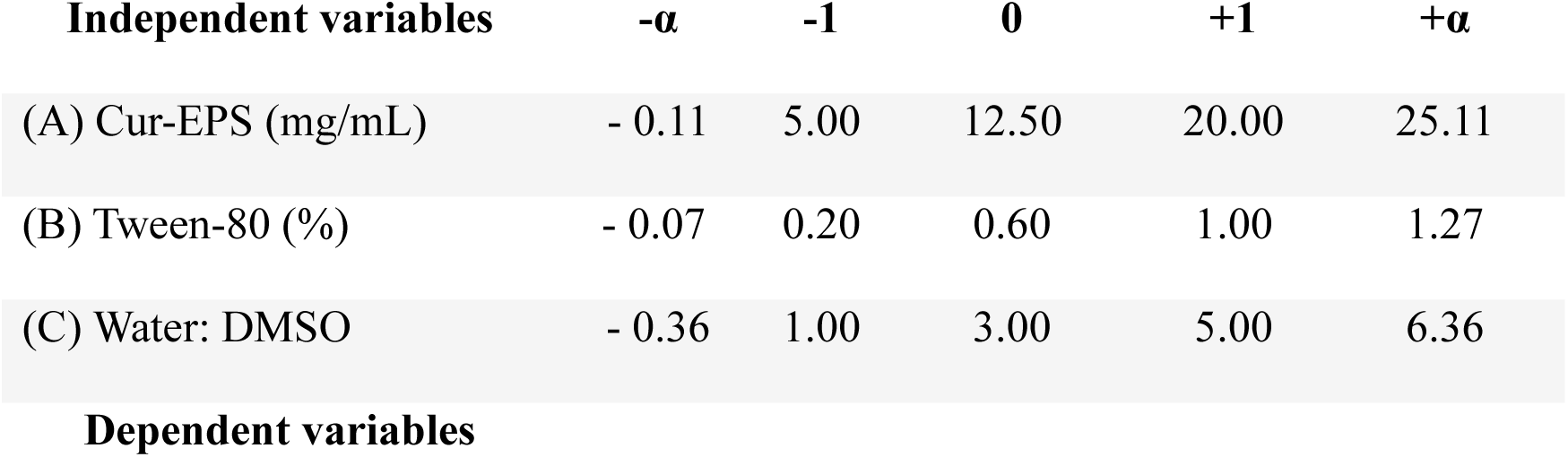
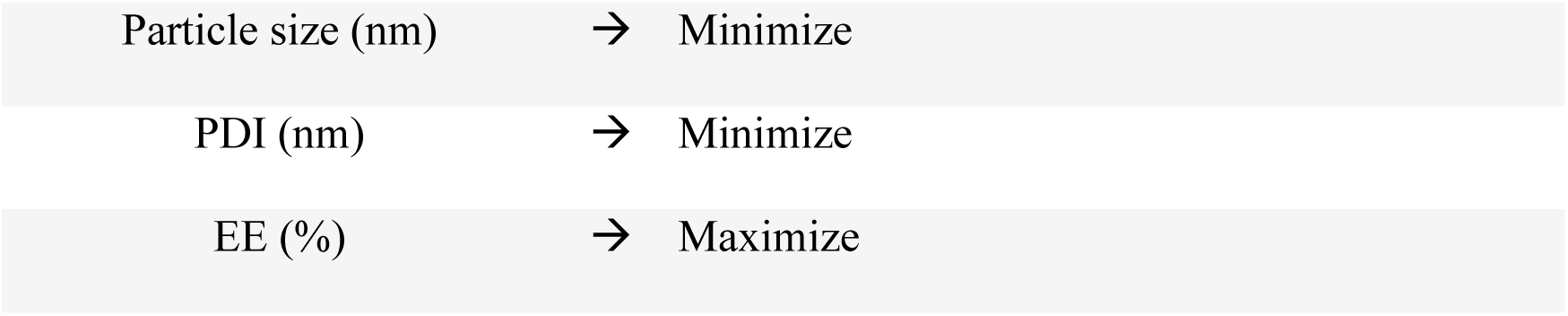
Variables and levels of design of experiments (DoE) for independent variables.

| Independent variables | -α | -1 | 0 | +1 | +$\alpha$ |
| --- | --- | --- | --- | --- | --- |
| (A) Cur-EPS (mg/mL) | - 0.11 | 5.00 | 12.50 | 20.00 | 25.11 |
| (B) Tween-80 (%) | - 0.07 | 0.20 | 0.60 | 1.00 | 1.27 |
| (C) Water: DMSO | - 0.36 | 1.00 | 3.00 | 5.00 | 6.36 |

**Dependent variables**
|  |  |  |
| --- | --- | --- |
| Particle size (nm) | → | Minimize |
| PDI (nm) | → | Minimize |
| EE (%) | → | Maximize |

The concentration of Cip was consistently maintained at 1 mg/mL throughout the entire experiment. The optimum combination of independent variables was chosen based on the criteria of minimizing the size and polydispersity index (PDI) range, while maximizing the entrapment efficiency. Finally, most efficient formulation was selected for the fabrication of Cip-Cur-EPS nanoparticle

### 2.4 Characterization of Cip loaded Cur-EPS

#### 2.4.1. Encapsulation and loading of Cur-EPS micelles with Cip

To determine the amount of Cip loaded on the micelle, the lyophilized Cip-Cur-EPS power was dispersed in Milli-Q (containing 0.05 N HCl) at concentration of 1mg/mL and was subjected to sonication for an hour. The sonicated solution was further centrifuged at 10,000 rpm for 15 min and the supernatant was collected in order to determine its absorbance at 275 nm with UV-visible spectrometer (Cary 60, Agilent Technologies, USA). The Cip concentration was determined by comparing absorbance to a standard calibration curve that was prepared by employing Cip solutions of 0, 2, 4, 6, 8 and10 μg/mL in Milli-Q water (Figure S). The baseline was established by measuring the absorbance of Cur-EPS micelles to verify that the micelles do not disrupt the absorbance measurements. The loading content (LC) and encapsulation efficiency (EE) were calculated using the following equations:

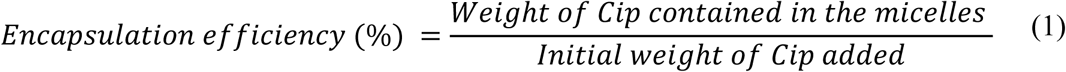

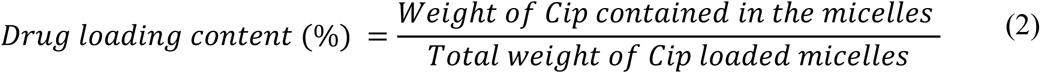

The yield of Cip-Cur-EPS recovered after the encapsulation process, was assessed by measuring the weight after freeze drying the suspension until a consistent weight was achieved.

#### 2.4.2. Particle size, PDI, zeta potential and morphology of Cip-Cur-EPS

The samples were diluted 1000 times in deionized water and were subjected to sonication using a bath sonicator (OSCAR Ultrasonic, Mumbai, India) for 5 min to obtain well dispersed suspension. The samples were quantified for their size using Litesizer (Anton Paar Litesizer 500, Austria) with scattering angle of 173°. Kalliope software was used to determine the hydrodynamic size, PDI and zeta potential. Measurements for all the samples were carried out in triplicate. The size of Cip encapsulated Cur-EPS was further analysed using Transmission Electron Microscopy (TEM, JEOL 1011, Japan). The Cip-Cur-EPS were prepared by depositing a micellar solution (0.1 mg/mL) on a carbon coated copper grid. The grid was desiccated at room temperature and subsequently examined using transmission electron microscopy (TEM) operated at 200 kV. The morphologies of Cip-Cur-EPS micelles were determined using a Field Emission Scanning Electron Microscope (FESEM, MERLIN, ZEISS, Germany). The lyophilized samples coated as thin film on an aluminium foil piece of 1 x 1 cm, which was further sputtered with gold and palladium and was subsequently examined under an accelerating voltage of 5.0 kV.

#### 2.4.3. Fourier transform infrared (FT-IR) spectroscopy

FT-IR analysis was performed to examine the interactions between Cip encapsulated within Cur-EPS micelles. Samples (Cip, Cur-EPS and Cip-Cur-EPS) in powdered form were compressed along with dry KBr powder into a disc for spectral scanning from wavenumber ranging from 450 to 4000 cm-1 by FT-IR spectrometer (Thermo Fischer NICOLET 6700, USA) and OMNIC 3.2 software.

#### 2.4.4. X-Ray diffraction (XRD)

A crystallographic analysis instrument X-ray diffractometer (XRD) (BRUKER D2 PHASER, USA) was utilized to ascertain the crystal-phase composition of the samples. The X-ray diffraction was monitored at 30 kV, Cu Kα radiation (γ = 1.54 Å) and 30 mA current, with scan speed of 2.0°/min and 2θ data collection range of 10° to 50°.

#### 2.4.5. *In-vitro* drug release

The release pattern of Cip from Cur-EPS micelles was investigated at different physiological pH (5.6 and 7.4) employing the dialysis method. Release of free Cip was also determined as control. The micelles, both empty and loaded with Cip (1mg/mL), were loaded inside a dialysis bag (MWCO-3,500). They were subsequently dialyzed against PBS or acetate buffer with pH values of 7.4 and 5.6 respectively, at 37 °C while being stirred at 50 rpm. Aliquots (2 mL) of the dialysis solution were collected at 30 min interval for 100 h. To maintain a constant volume of the stock dialysis solution, 2 mL of fresh buffer was added after each aliquot removed. The absorbance of the Cip released in the buffers of different pH were assessed using a UV-visible spectrophotometer at 275 nm. By utilizing the calibration curve of Cip established across a concentration range of 0 to 10 μg/mL, the UV absorbance was correlated to the amount of Cip released from the micelles. The cumulative drug release was determined using the following equation:

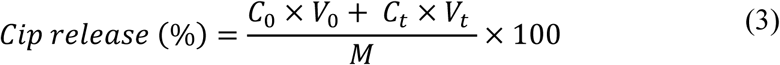

where *V_o_* is the total volume of release buffer (bath volume), *C_o_* is the concentration of Cip in the bath volume, *V_t_* is the volume of the release buffer removed at a specific time *t*, *C_t_* is the concentration of Cip released at a specific time, *M* is the concentration of Cip loaded within the Cur-EPS micelles.

#### 2.4.6. Release kinetics models

Several drug release kinetic models, such as zero-order kinetics, first-order kinetics, Higuchi model, Hixson-Crowell model and Korsmeyer-Peppas model were employed to evaluate the release kinetics of Cip. Further details on the release kinetics are presented in the supplementary section (Appendix S1).

#### 2.4.7. Storage stability of Cip-Cur-EPS

The Cip-Cur-EPS in its solution form in PBS was determined at 4 °C for a period of 14 days. The stability of Cip-Cur-EPS solution was monitored after every 2 days by measuring the hydrodynamic size (nm), PDI and EE (%) as described in the previous section.

### 2.5. *In vitro* antibiofilm activity

The blank Cur-EPS micelles, Cip and Cip-Cur-EPS were assessed for their antibiofilm activity against MRSA (Methicillin-resistant *Staphylococcus aureus*) (ATCC 33591) and *P. aeruginosa* (ATCC 27853). More details on the methods to determine the *in vitro* antibiofilm activity and the time-kill kinetics for the disruption of mature biofilms were presented in supplementary information (Appendix S2).

### 2.6. Visualization of microbial biofilm removal by confocal laser scanning microscopy (CLSM)

The bacterial cultures (1×10^6^ CFU/mL) were added into a 24 well plate with sterile round glass coverslips (18 mm diameter) after being cultured overnight. The wells were filled with TSB medium and the biofilm was allowed to develop for 72 h at 37°C under static condition. The slides were thoroughly rinsed with PBS three times in order to eliminate any bacteria that did not adhere to the surface. 1 mL of fresh medium, supplemented with the test samples were added in the wells. The plates were incubated at 37 °C for 72 h. Cells cultured in a medium without any samples were considered as control. Ultimately, the slides were rinsed thrice with PBS and thereafter treated with 100 µL of acridine orange (AO) (0.01%, w/v) for a duration of 10 min at room temperature in the absence of light. In addition, the coverslips that had been stained were carefully rinsed using PBS and were then examined using CLSM (Olympus FluoView FV3000, Japan). The stained biofilm observed by employing excitation wavelengths of 488 nm argon laser and the emitted fluorescence was detected with a long-pass filter at 500−535 nm. The images were obtained from several locations of biofilms that developed on the upper surface of the coverslips. The images were observed using Olympus FluoView and ImageJ software was employed to gather z-stacks for three-dimensional (3D) reconstruction of biofilm images. The experiments were performed in duplicate, and a minimum of 5 fields were collected for each sample. The method for scanning electron microscopy (SEM) analysis of biofilms is mentioned in the supplementary section (Appendix S3).

### 2.7. Quantitative analysis of biofilm structure

COMSTAT analysis was conducted using data acquired from the green channel used to visualize AO-stained biofilms. The images of the stacks obtained from CLSM were exported from the Olympus FlourView Image software in form of raw series and were converted into grey scale and were saved as tagged image file format (TIFF) with Image J software. Further, COMSTAT 2 (version 2.1) was employed for quantitative analysis of biofilm. Several COMSTAT function were employed to determine the mean thickness of the biofilm (µm) and the biovolume of each image stack is expressed as the volume of biomass per substratum area (µm^3^/µm^2^) that evaluates variability in the biofilm thickness as well as the structural heterogeneity.

### 2.8. Maintenance of cultures in the Artificial urine medium (AUM)

The protocol for preparation of AUM and preliminary studies to determine the growth of *P. aeruginosa* and MRSA were included in the supplementary section (Appendix S4).

### 2.9. *In-vitro* eradication of *P. aeruginosa* and MRSA biofilm formed in the urinary catheter

Foley catheters (6mmØ, 16 cm length) (Haiyan Kangyuan Medical Inst Co., Ltd. Zhejiang, China), a category of indwelling catheters intended for temporary or prolonged bladder retention, were sectioned into 1 cm segments and sterilized in 70 % ethanol for 1 h and were allowed to dry under a flow cabinet for 30 min. The sterilized catheter pieces were placed in 35 mm cell culture petri plates (Tarsons, South Korea) containing AUM (2 mL) were inoculated with bacterial culture (200 μL, 10^6^ CFU/mL) and were incubated for 72 h at 37 °C. Subsequently, the AUM was removed from petri plates and subsequently replaced with fresh AUM supplemented with the test samples. The catheter pieces placed in AUM without any test samples were included as controls. Following the incubation period for 72 h, the catheter pieces were aseptically removed and were washed with 0.9 % saline solution to remove planktonic bacterial cells, and subsequently placed into sterile Eppendorf tubes containing 1mL of 0.9 % saline solution. The tubes were vortexed for 5 min followed by sonication in for 1 min. The sonicated suspensions were serially diluted in 0.9 % saline solution and were plated on TSA, which were further incubated for 24 h at 37 °C to determine the viable bacteria after antimicrobial treatment in terms of colony forming units in the catheter pieces. The log reduction values (LRV) were calculated using following formula to express the relative number of living microbes that are eliminated by treatment with the samples:

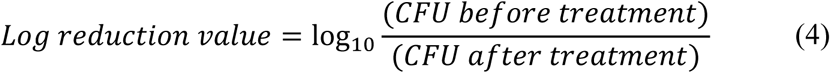

The biofilm biomass was determined by Crystal Violet (CV) staining. The catheter pieces were removed and washed with PBS and were further transferred into 35 mm plate containing 0.1% CV and were incubated at 37 °C for 15 min. The samples were rinsed with saline solution and were further air-dried. The remaining CV on the catheter pieces were dissolved in 95% ethanol. 200 μl from each plate were transferred into a 96-well plate and the optical density of each well was measured at 570 nm, using a microplate reader (Thermoscientific Multiskan Go, USA). The data were presented in form of biofilm inhibition percentage as mentioned earlier.

#### 2.9.1. Effectiveness of treatments in suppressing the bacterial re-growth

Treatment with blank Cur-EPS micelle, free Cip and Cip loaded Cur-EPS micelle to prevent the re-growth of MRSA and *P. aeruginosa* on the catheter was also determined. (16) The catheter pieces treated with samples as described above, were immersed in 5 ml of fresh TSB, which were further re-incubated for 24 h at 37 °C. After incubation period, the catheter pieces were washed with saline solution and the bacterial cells were recovered by sonication as described in the above section. The bacterial suspensions were used to determine live bacterial cells from each catheter piece by determining colony forming units on the agar plates.

#### 2.9.2. SEM of the urinary catheter pieces

The ability of Cip, in both its free and loaded form within Cur-EPS, to eradicate biofilms formed by MRSA and *P. aeruginosa* on urinary catheters was examined via a scanning electron microscope, following previously established method. (17) The catheters were cut in pieces along the cross section and then washed three times with PBS to eliminate any unattached cells. Subsequently, the samples were moved to a 24 well plate and the bacterial cells were fixed with 2.5% glutaraldehyde at 4°C for 12 h. Subsequently, the catheter pieces were rinsed with PBS and dehydrated using varying concentrations of ethanol (50%, 70%, 80%, 90%, 95%, and 100%). The catheter pieces with attached biofilm were subjected to freeze-drying and thereafter stored in a desiccator. The specimens were secured onto aluminium stubs using carbon tabs and colloidal silver paste. They were then coated with a layer of gold-palladium using a Hummer VI sputter coater (Anatech in Battle Creek, Michigan, USA). The samples were observed using a Quanta 200 scanning electron microscope (FEI, Hillsboro, Oregon, USA) with an operating voltage of 25 kV.

### 2.10. *In Vitro* Model of Catheterized Bladder

*In vitro* model of catheterized bladder was used to assess the ability blank Cur-EPS, Cip, Cip-Cur-EPS to prevent biofilm formation on the Foley urinary catheters under dynamic settings. The catheter was inserted in an *in vitro* model of the human bladder and was constantly supplied with AUM using a peristaltic pump (Watson Marlow, Devens, USA) at a flow rate of 0.5 mL/min. This flow rate was chosen to account for the rate of glomerular filtration and reabsorption of primary urine in the renal tubules. The catheter was inserted into the bladder model and fixed by inflation of the balloon with 5 mL of PBS. The bladder vessel was then filled till the catheter eye with sterile AUM supplemented with the bacterial culture (10^6^ CFU/mL) in AUM. The model was maintained for 5 days inside an incubator at 37 °C. Subsequently AUM supplemented with the test samples were introduced into the catheters, which were further incubated for 72 h. The catheters were removed and the tubes were gently flushed with PBS. Subsequently, the tubes were filled with a crystal violet solution (0.1%) and were incubated for 20 min at room temperature to stain the biofilm followed by another rise with PBS. The tubes were visually photographed in order to evaluate the dyed biofilm. Furthermore, the tubes were filled with 95 % ethanol and the liquid was collected to measure the absorbance at 570 nm to determine the biofilm biomass. The data were presented in terms of biofilm eradication percentage.

### 2.11. Statistical Analysis

All the experiments were conducted in triplicate, and the findings were expressed as mean ± standard deviation (SD). To compare the results of each variable, Student’s t-test and analysis of variance (ANOVA) were employed, followed by Tukey’s post-hoc test, with the *p*-value < 0.05, *p*-value < 0.01 and *p*-value < 0.001 being indicative of the statistically significant difference.

## 3. Results and discussion

### 3.1. Optimization of Cip-Cur-EPS micelles prepared by nanoprecipitation method

The Cip was loaded within Cur-EPS micelles via nanoprecipitation (Figure 1). This approach relies on the differences in surface tension between the aqueous phase and the water-miscible organic solvent phase. When the two phases are mixed, the variation in the surface tension and the mutual miscibility between the phases leads to the migration of the organic solvent phase towards the aqueous phase causing the precipitation of the particles, which culminates in the generation of nanoparticles at the interphase. Nanoprecipitation has been utilized for incorporating hydrophobic bioactive compounds, including quercetin, paclitaxel, cucurbitacin, and amphotericin B etc., into colloidal drug delivery systems. This approach has gained significance in pharmaceutical product development mainly due to the use of low-toxicity solvents, simple procedure, and minimal energy consumption. (19)

**Figure 1.**
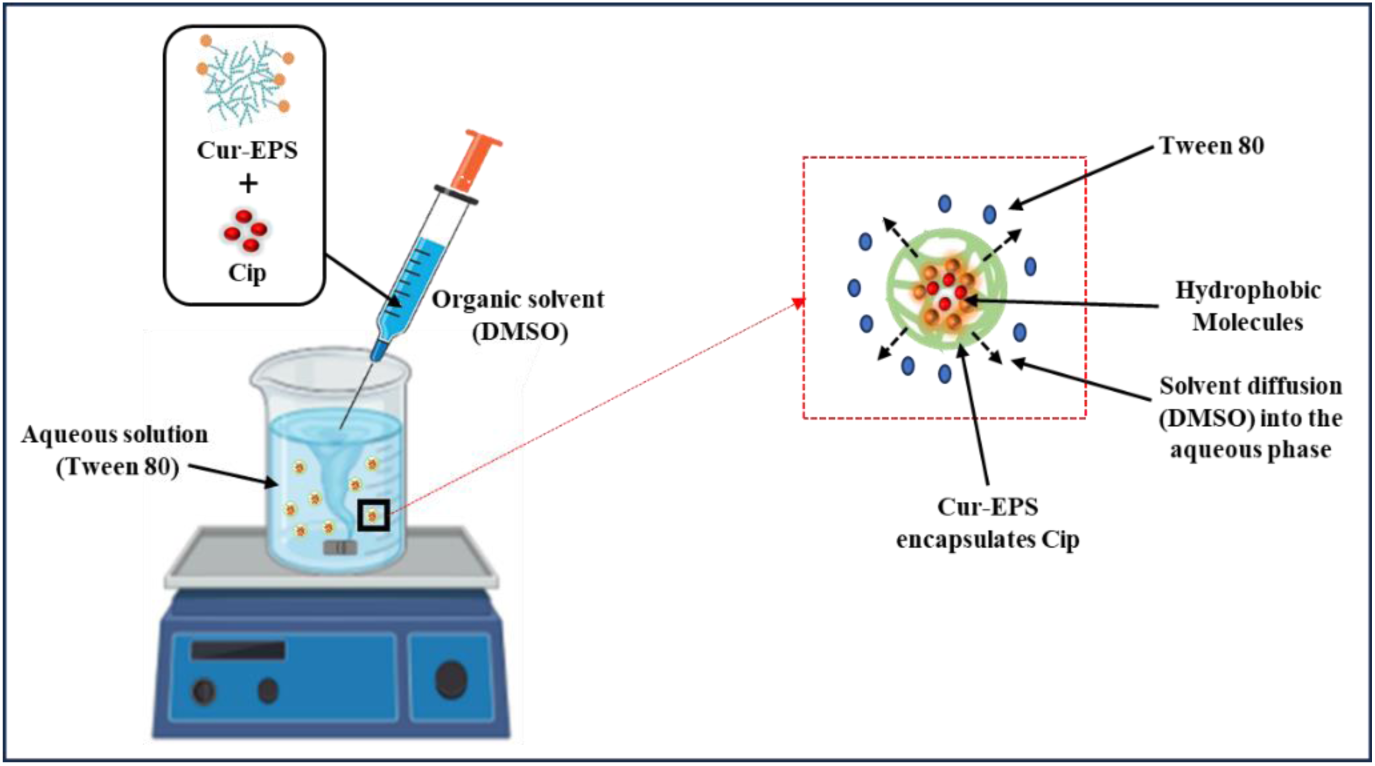
Preparation of Cip-Cur-EPS by nanoprecipitation method. (18)

The main goal of this study was to evaluate the effect of Cip loaded within Cur-EPS micelles to eradicate the bacterial biofilms. Thus, the formulation of Cip-Cur-EPS was optimized to enhance the encapsulation of the Cip within the Cur-EPS micelles as well as to enable homogeneous distribution and penetration of the micelles within the biofilms in order to achieve optimal biofilm eradication effect. Therefore, this study employs RSM to determine the interactions of several factors such as Cur-EPS concentration (A), Tween 80 concentration (B) and water: DMSO ratio (C) in elucidating their impact on the main characteristics of Cip loaded Cur-EPS micelles such as particle size (X1), PDI (X2) and EE (X3). The predicted and the experimental results related to the assessed factors on the responses to fabricate Cip loaded Cur-EPS micelles are presented in Table S1. The three responses from the 20 runs exhibited considerable variability, with values of particle size, PDI and EE ranged from 304.34 nm to 392.91 nm, 0.38 to 0.69 and 63.03 % to 86.97 % respectively. A mathematical relationship between factors and responses was established by response surface regression analysis utilizing Minitab® 22.1 software. The significance of the model’s coefficients was determined from the *F* and *p* values from the ANOVA results (Table S2, S3 and S4). For each response, the model which generated high *F* value and *p*-value < 0.05 were identified as the best fitted model. The responses X1, X2 and X3 were significantly influenced by factor A (*F* value = 958.82), factor B (*F* value = 229.11), and factor C (*F* value = 295.20), respectively. Furthermore, the impact of all linear variables, including A, B, and C, were observed to be highly significant on the response X1 and X3 (*p* values < 0.05). Nonetheless, for X2, all linear variables, excluding factor A (*p* > 0.05), were deemed significant. The quadratic terms of A, B, and C (A^2^, B^2^, and C^2^) had substantial effects (*p* < 0.05) on all three responses. The interaction effects of AB affected X1 and X3 (*p* < 0.05). The interaction effects AC and BC had no impact on any of the observed responses (*p* > 0.05).

The regression equations for three responses were expressed in terms of coded factors have been presented in the following equations.

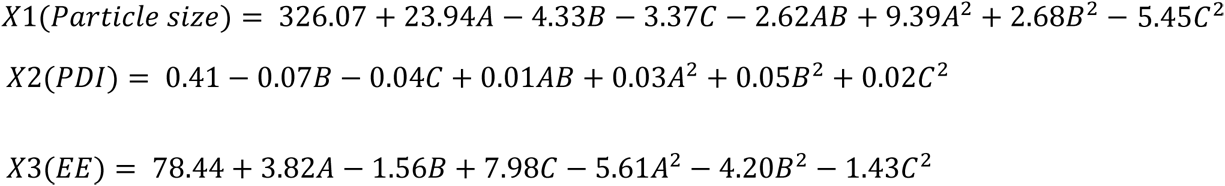

In a regression equation for a response, a positive value signifies a synergistic effect whereas a negative value denotes an inverse association between the factors and the response. (20) The regression equation for X1 indicated that factor A, A^2^ and B^2^ exerted a synergistic effect. Moreover, X1 was also affected by the antagonistic effect of B, C, C^2^ and AB. X2 was influenced by the synergistic effect of A^2^, B^2^, C^2^ and AB and the antagonistic effects of factors B and C. The X3 was impacted by the synergistic effect of A and C and antagonistic effect of B, A^2^, B^2^ and C^2^. The models for all the three responses demonstrated a good conformity with experimental data, as evidenced by high Adj. R^2^ values (Table S2, S3 and S4). The closer the value of Adj. R^2^ to R^2^, the greater is the power of model to predict responses. Adj R^2^ and R^2^ were found to be in agreement with each other. Also, the Pred.R^2^ values (0.95 for particle size, 0.94 for PDI and EE) were in a satisfactory agreement with Adj.R^2^ values (0.98 for particle size, 0.96 for PDI and 0.97 for EE). The lack of fit was not statistically significant at the 95% confidence level.

Furthermore, the normality of the data can be assessed by generating the NPP (normal probability plot) of the residuals. The NPP is a graphical method for evaluating the approximate normality of a data set. The residual is the variation between the observed value and the predicted value derived from the regression analysis. The data are normally distributed if the points on the plot are closely aligned with the straight line. The Figure S1. illustrates the normal probability plot of residual values. The experimental points exhibited a reasonable alignment, indicating a normal distribution.

The main effects and the interactive effects of two factors on the response in the RSM were further analysed using 3D response surface graphs and contour plots (represents the response surface as a 2D projection) (Figure 2) The response surface plots indicated that the particle size and EE increased with elevated concentration of Cur-EPS. An increase in the polymer concentration often enhanced particle size, PDI and EE. Previous reports also indicated that an increase in the polymer concentration resulted in a rise of particle size and encapsulation efficiency for the optimization of Ansamycin-loaded PLGA nanoparticles. (21) This may be attributed to an increase in the viscosity of the organic solvent phase (DMSO), resulting in larger particles with multimodal distribution and impeding the diffusion of the drug from organic to aqueous phase. An increase in the amount of Tween 80 led to the reduction in particle size and PDI. A rise in surfactant concentration improves the interfacial stability, which leads to a reduction of particle size and PDI. (22) However, a significant increase in the amount of Tween 80 resulted in the decline in EE. Elevated surfactant concentration in the external aqueous phase may facilitate the diffusion of drugs from the organic solvent phase and eventually they might be soluble in the aqueous medium in form of micelles thereby reducing the EE. Additionally, the significant increase surfactant concentration can elevate the viscosity of external aqueous phase thereby hampering the effective diffusion of organic phase leading to larger droplet formation and thus resulting in the increase of the particle size. (22)

**Figure 2.**
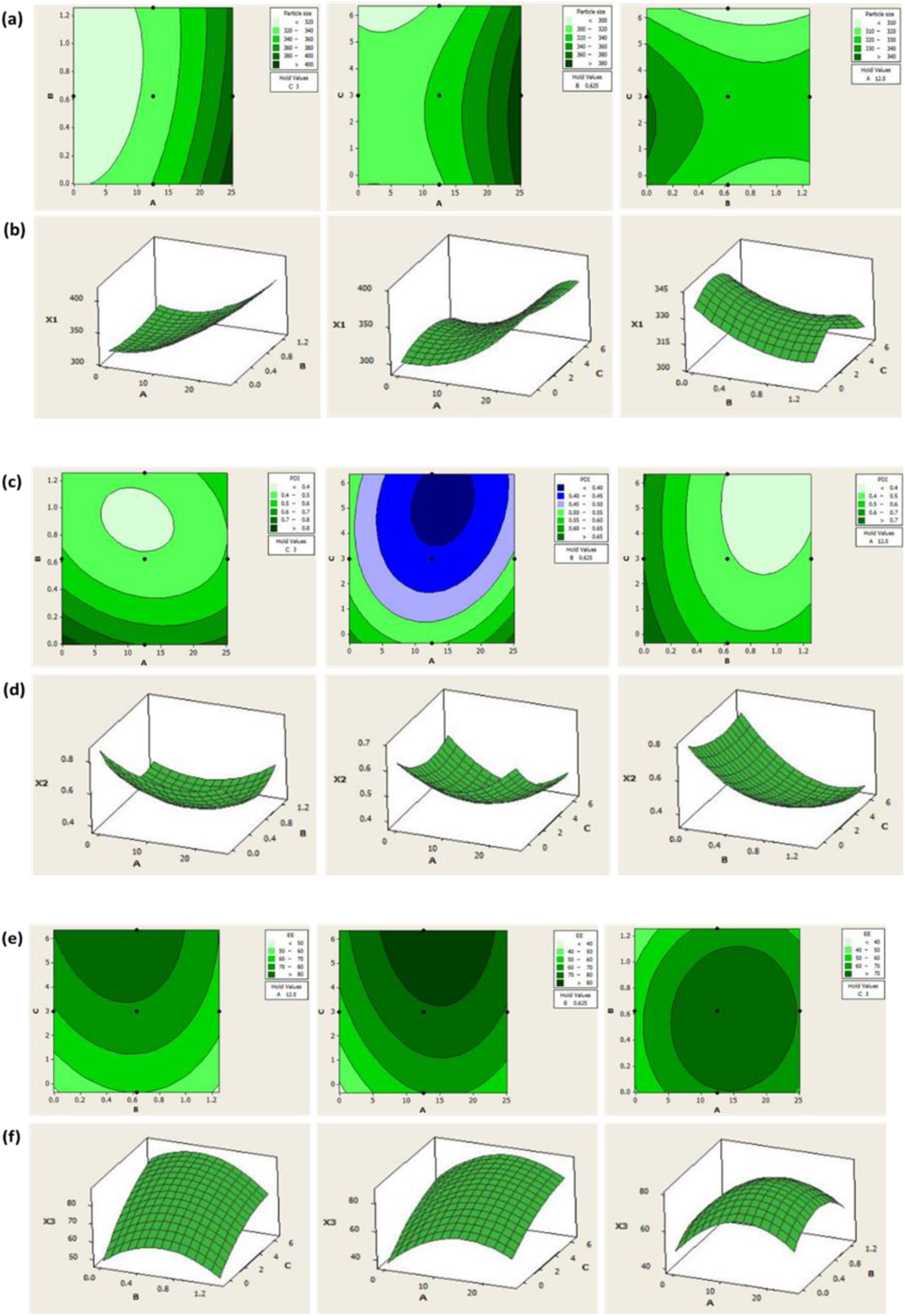
Contour plots (a) and 3D response surface plots (b) for particle size (X1), contour plots (c) and 3D response surface plots (d) for PDI (X2) and contour plots (e) and 3D response surface plots (f) for EE (X3). The factors are denoted as A, B and C which indicates Cur-EPS concentration (mg/mL), Tween 80 (%) and water: DMSO respectively.

Furthermore, an increase in the aqueous to organic phase ratio results in a reduction of particle size and PDI, while simultaneously enhancing the EE. In nanoprecipitation method the nanoparticles are formed as a result of quick solvent diffusion into the aqueous phase. Consequently, as the volume of the aqueous phase escalates, the diffusion of the organic solvent inside the aqueous phase increases, resulting in a reduction of particle size while improving the particle homogeneity and EE. However, a further increase in the volume of the aqueous phase may enhance the amount of drug that can be solubilized in water, resulting in greater drug loss into the aqueous phase.

The particle size is a crucial determinant in nanoparticle-based drug delivery systems. A particle size of < 500 nm is advised for effective biofilm management since particle with larger sizes can obstruct biofilm infiltration and also affect its systemic circulation stimulate the complement system, leading to rapid identification of such particles by the immune system leading to subsequent elimination from the bloodstream. (9) The PDI another essential parameter used to characterize the variance in particle size within a sample and an ideal value is preferred to be nearer to 0. Finally, high EE is desirable since it delivers adequate amount of drug to the target site and prolongs the drug’s residence time. Therefore, the main goal of this study is to develop a formulation for Cip-loaded Cur-EPS micelles featuring smaller particle size, a low PDI, and high EE that exhibits robust antibiofilm activity. However, optimizing all the responses together is challenging due to their lack of alignment. The ideal condition achieved in one response may adversely affect another response. The multi-criteria problem can be addressed as a single criterion by employing the global desirability function (D) to identify the optimal formulation for all responses. (20) For concurrent optimization, each response was assigned a low and high value. The EE was adjusted to maximize whereas the particle size and PDI were aimed to attain a minimal goal. Consequently, individual desirability function was obtained for each response, with values represented on a non-dimensional scale spanning the interval 0 – 1. A maximum D value of 0.96 was attained at optimal concentrations of the independent variables (Figure S2). Furthermore, Cip-Cur-EPS was prepared utilizing the optimal independent variables to assess the prediction capability of the model. The experimental outcomes were compared with the model-predicted values for all three responses (Table 2). The experimental values obtained under ideal conditions closely aligned with the predicted values, exhibiting low percentage bias, indicating that the optimized formulation was reliable. Therefore, high EE, alongside a minimal particle size and particle size distribution were achieved by employing optimal formulation factors (Cur-EPS - 11.06 mg; tween 80 - 0.72 %; water : DMSO - 6.36).

**Table 2.** The predicted results acquired by RSM and the empirical data for the corresponding responses under the optimal conditions for the formulation of Cip-Cur-EPS.

| **Optimized factors** | **Cur-EPS concentration** | **Tween 80 concentration** | **Water : DMSO** |
| --- | --- | --- | --- |
|  | 11.863 | 0.74 | 6.36 |
| **Responses** | **Particle size** | **PDI** | **Encapsulation efficiency (%)** |
| **Predicted** | 300.05 | 0.37 | 85.07 |
| **Experimental** | 303.82 | 0.36 | 84.95 |
| **Bias (%)** | -1.25 | 2.70 | 1.18 |
| Bias was calculated as (predicted value – experimental value) / predicted value × 100<br>(Acceptance criteria 6%) |  |  |  |

### 3.2. Characterization of Cip-Cur-EPS micelles

The characteristics of the optimized Cip-loaded micelle were presented in Table 3. The incorporation of Cip within Cur-EPS micelles via nanoprecipitation was confirmed by assessing the Cip encapsulation efficiency and loading content, which were determined to be 84.50 ± 2.01 % and 9.41 ± 0.39 %, respectively (Table 2) with the loaded Cip concentration of 90 ± 3.11 µg/mL. A favourable Cip-Cur-EPS yield of 78.63 ± 2.49 % was attained. In prior studies, the high concentration of Cip (>1 mg/mL) was used to enhance the drug loading capacity. (23, 24) Treatment with high dosage of antibiotics often leads to antimicrobial resistance. In this study, the initial Cip concentration was maintained at 1 mg/mL throughout the formulation process, as the goal this study was to develop Cip loaded carrier with low Cip concentration while attaining good antibiofilm effect. Such formulations can improve the therapeutic index of the Cip while reducing its side effects for treating biofilm infections.

**Table 3.**
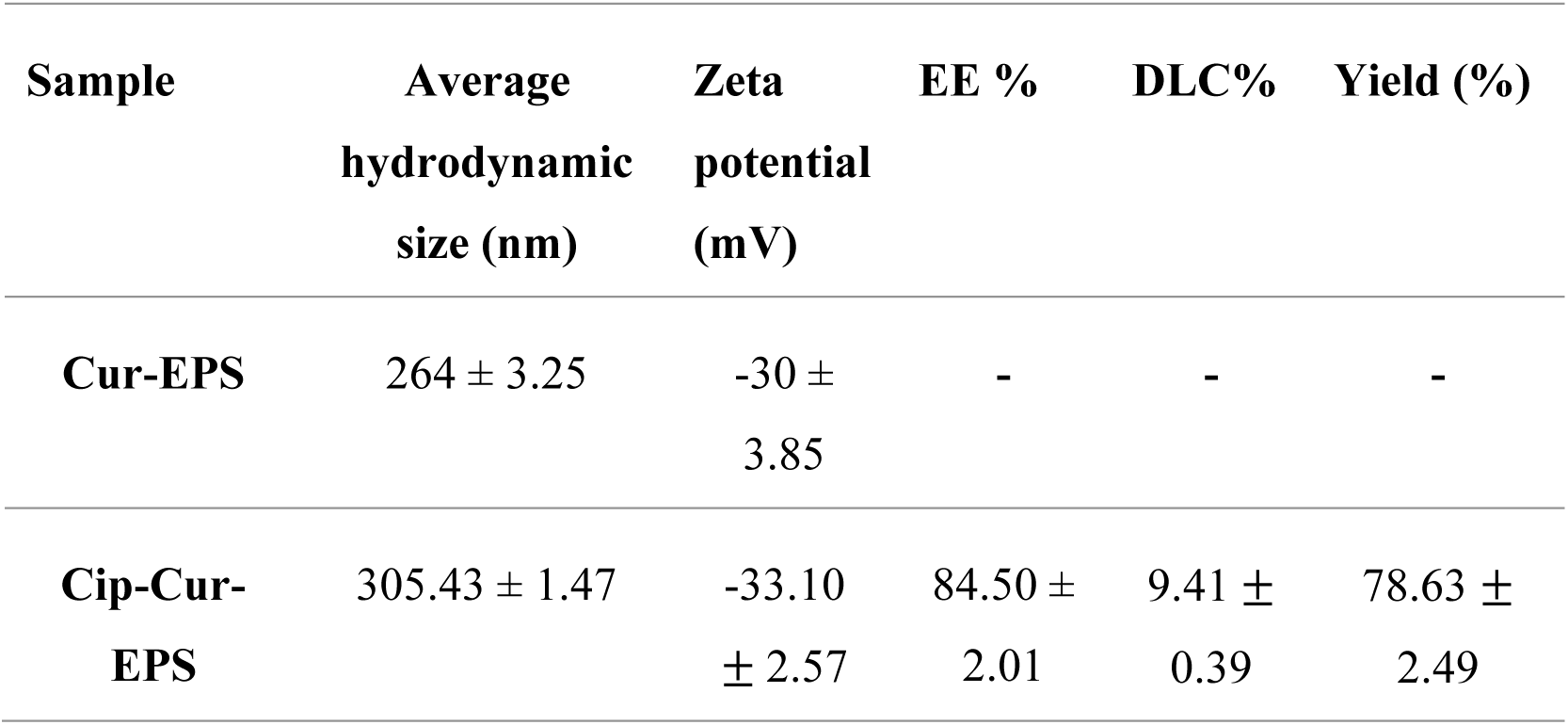
Characteristics of Cip-Cur-EPS micelles in terms of average hydrodynamic size, zeta potential, EE, DLC and yield.

| Sample | Average hydrodynamic size (nm) | Zeta potential (mV) | EE % | DLC% | Yield (%) |
| --- | --- | --- | --- | --- | --- |
| Cur-EPS | 264 ± 3.25 | -30 ± 3.85 | - | - | - |
| Cip-Cur-EPS | 305.43 ± 1.47 | -33.10 ± 2.57 | 84.50 ± 2.01 | 9.41 ± 0.39 | 78.63 ± 2.49 |

Furthermore, the improvement in the solubility of Cip within Cip-Cur-EPS was also observed. At physiological pH, the aqueous solubility of Cip is significantly affected due the transfer of protons from the carboxylic acid to the piperazine ring, which is a basic component, results in the formation of zwitterionic species. Cip is almost insoluble at physiological pH, exhibiting a hydrophilicity index log P < 0.1, which causes drug precipitation before it reaches the afflicted area, hence resulting in limited bioavailability. (25) Moreover, the incorporation of Cip within Cur-EPS micelles considerably enhanced the solubility of Cip in PBS compared to its free form, which precipitated in PBS, as illustrated in Figure 3A. The Cip was solubilized by the Cur-EPS micelles through the hydrophobic interactions with the micellar core.

**Figure 3.**
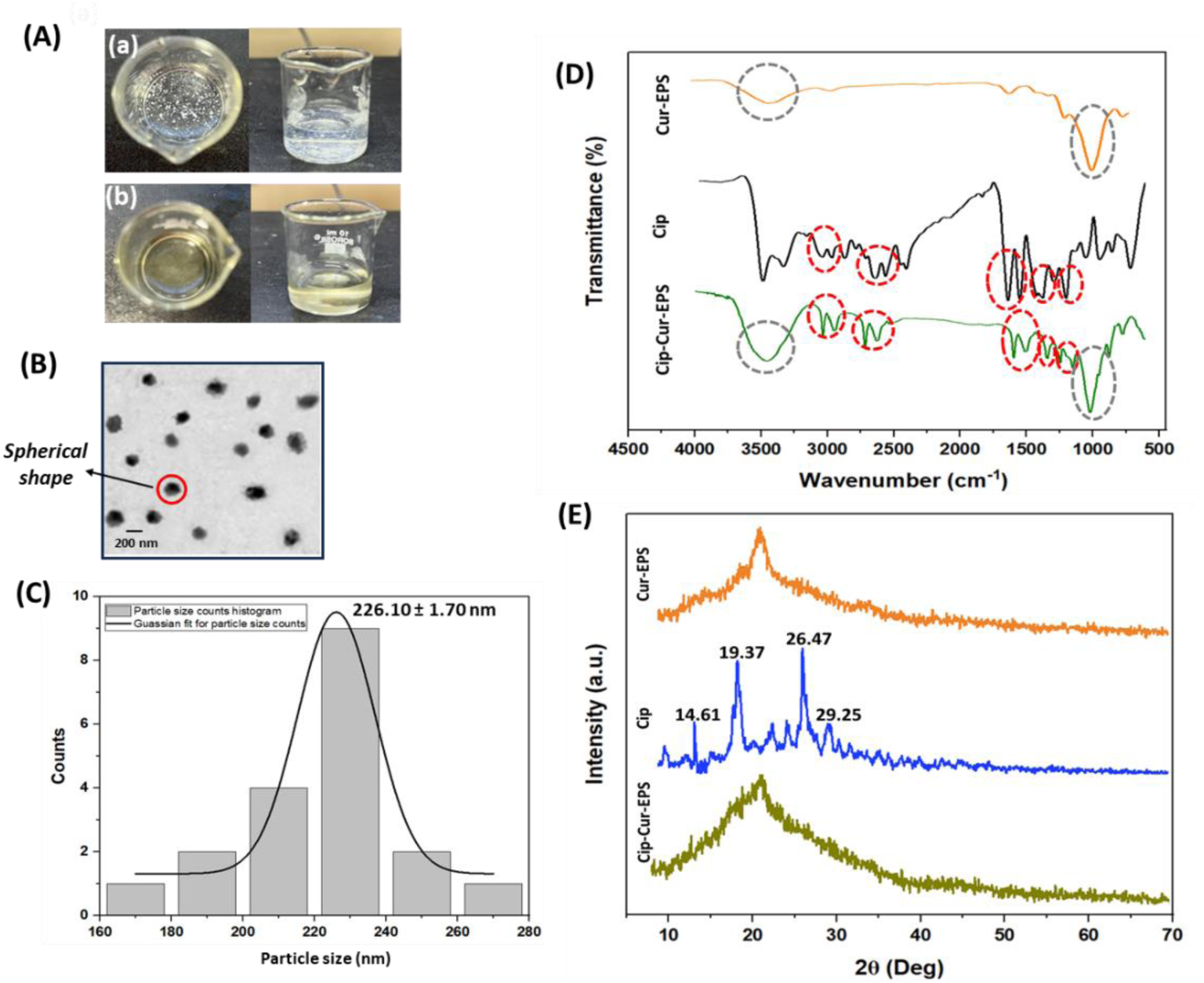
(A) Solubility of Cip in PBS in (a) free form and (b) Loaded form in Cur-EPS; (B) TEM image (50 K X magnification); (C) Size distribution histogram based on the TEM image (Image J software) of Cip loaded Cur-EPS micelles; (D) FT-IR spectrum of blank Cur-EPS micelles, Cip and Cip loaded Cur-EPS micelles. Cip-Cur-EPS micelles exhibited the distinctive peaks of Cip (represented by red circles) and the peaks related to Cur-EPS micelle (represented by grey circles); XRD patterns of blank Cur-EPS micelles, Cip and Cip loaded Cur-EPS micelles.

The size and the PDI of the optimized Cip-Cur-EPS micelles were measured by DLS. As mentioned in the previous section Cip-Cur-EPS prepared by optimal formulation parameters exhibited average hydrodynamic size of 305.43 ± 1.47 nm. Cip occupied space within the hydrophobic core of the Cur-EPS micelles, which resulted in increase of the average hydrodynamic size compared to the blank micelle (254 ± 2.7 nm). (11) Moreover, the Cip-Cur-EPS exhibited relatively narrow size distributions with a PDI of 0.38 ± 1.23, which was favourable for particle stability.

However, the DLS measurements are limited to providing information solely on the hydrodynamic size of the particles, without offering significant insights into its shape. Therefore, TEM was performed to provide a more precise indication on the formation of Cip-Cur-EPS (Figure 3B). The TEM image and the corresponding histogram depicting the particle size distribution of Cip-CUR-EPS were presented in Figure 3C. The average size was found to be 226 ± 1.70 nm and the Cip loaded micelles were observed to exhibit spherical shape. The variations in size of the micelles, as determined by TEM and DLS, can be attributed to the fact that DLS measures the hydrodynamic diameter of the micelles in water, while samples used for electron microscopy underwent shrinkage as a result of water evaporation during air-drying. (26) Furthermore, FESEM image in the supplementary section (Figure S3) presents the surface morphology of Cip-Cur-EPS. The electron microscopy images affirms that the micelles loaded with Cip exhibited spherical morphology.

Furthermore, the surface charge of the Cip-Cur-EPS micelles in solution was assessed by determining the zeta potential. The zeta potential serves as a reliable measure of colloidal stability of the particles in the solution, with values above + 30 mV or falling below - 30 mV are regarded stable. (27) The surface charge of Cip-Cur-EPS micelles was measured to be - 33.10 ± 2.57 mV, suggesting their considerable stability under physiological conditions at pH-7.4.

FT-IR spectra of free Cip, blank Cur-EPS, and Cip-Cur-EPS were presented in Figure 3D. The FTIR spectrum of Cip exhibited multiple transmittance bands corresponding to specific molecular vibrations. These include the O−H stretching in COOH with the wavenumber range of 3530 cm^−1^ - 3379 cm^−1^, the stretching of symmetrical and asymmetrical stretching vibrations of C−H bonds within the wavenumber range of 3000 - 2900 cm^-1^, the N−H stretching within the range of 2650 - 2465 cm^−1^, the carbonyl group (C=O) stretching in carboxylic acid and N−H bend of pyridone moiety at 1708 cm^−1^ and 1620 cm^−1^, respectively. Additionally, the spectrum for Cip exhibited the C−N and the C−F stretching vibrations at 1375 cm^−1^ and 1272 cm^−1^ respectively. The FT-IR spectrum of Cip exhibited characteristic peaks, which were consistent with those reported in the literature. (28) Moreover, the blank Cur-EPS micelle showed bands including O−H stretching at 3500 cm^-1^, C−H aliphatic group at 2882 cm^-1^, C=O stretching at 1700 cm^-1^ and C−O−C bonds vibrations of the glycosidic bridges at 1000 cm^-1^ as reported previously. (11) The FTIR spectra of Cip loaded Cur-EPS micelles demonstrated the presence of the distinctive peaks related to both Cip and blank Cur-EPS micelles. There were no notable alterations in the position of the peaks of free Cip in the Cip-Cur-EPS, indicating that there were no significant chemical interactions between Cip and Cur-EPS micelles during the synthesis of Cip-loaded Cur-EPS micelles. These observations also suggested that there were no potential inconsistencies between Cip and Cur-EPS in Cip-Cur-EPS indicating the compatibility between the drug and the excipient. The results aligned with the previously reported literature, where Cip was encapsulated within xanthan gum - chitosan based carrier. (23)

Furthermore, the X-ray diffractograms patterns of Cip, Cur-EPS and Cip loaded Cur-EPS are presented in Figure 3E. The XRD patterns of Cip exhibited a distinct diffraction pattern at 2θ of 14.61, 19.37, 26.47, and 29.25 respectively. The sharp peaks within the 2θ range of 10°-40° confirmed the crystalline form of Cip. However, the resolution and accuracy of XRD became compromised in case of Cur-EPS micelle, as the majority of the EPS backbone in Cur-EPS exhibits significant amorphous features with diverse interatomic distances. (29) Consequently, Cip did not exhibit sharp peaks when loaded within Cur-EPS micelle. Therefore, the crystallinity of Cip disappeared within the Cur-EPS micelles indicating that the formulation was amorphous in nature. Thus, based on this observation, it can be deduced that Cip was loaded within Cur-EPS micelles. Earlier reports have also inferred that the crystallinity of encapsulated Cip was completely masked within the starch-based microspheres. (30) The amorphous form of a drug is crucial in therapeutic development due to several advantages compared to its crystalline counterpart, including increased free energy, structural disorder, enhanced water solubility, and molecular mobility. (31) Therefore, the amorphous form of Cip-Cur-EPS can enhance the solubility of Cip under physiological conditions, thereby increasing its therapeutic value.

### 3.3. *In-vitro* release of Cip from Cur-EPS micelles

This study examined the utilization of Cur-EPS micelles as an antibiotic carrier, featuring pH-responsive release of the loaded antibiotic at physiological temperature (37 °C) to eradicate robust bacterial biofilms. The release of the Cip loaded within Cur-EPS micelles were studied in the phosphate buffer with pH 7.4, which mimics the pH of extracellular fluid, and in an acetate buffer with pH 5.6, which resembles the pH found at a biofilm infection site. Figure 4 displays the Cip release from the micelles at different pH (5.6 and 7.4). A preliminary burst release is necessary for the delivery of antibiotics to limit bacterial growth at the initial phase; but, for persistent bacterial populations, such as those present in biofilms, a sustained release of antibiotics may be advantageous to eradicate them. Therefore, Cip-Cur-EPS can be effective in mitigating biofilms as the Cip release profiles can be categorized into two stages i.e., the initial immediate release of Cip was detected within the first 4 h, followed by the sustained release of Cip that appeared for up to 60 h at pH 5.6 and for up to 30 h at pH 7.4. The Cip-Cur-EPS formulation released 50 % of Cip within 30 h at pH 5.6. Subsequently, the rate of release decelerates and stabilizes after 30 h and 60 h for pH levels of 7.4 and 5.6 respectively. The influence of the pH on the rate of release becomes insignificant at this juncture.

**Figure 4.**
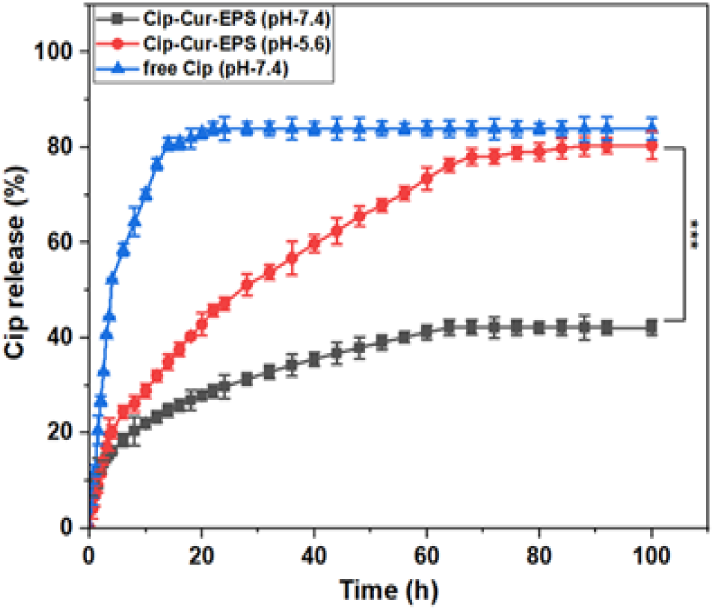
(A) Release profiles of free Cip and Cip from Cur-EPS in acetate buffer (pH - 5.6) and PBS (pH - 7.4). Data is presented as mean values and standard deviation of three independent experiments. Significant levels indicated by asterisks (*\*\*\* p < 0.001*).

Comparable findings concerning the release of Cip were also reported earlier. (32) Furthermore, at pH 5.6, of the Cip release was significantly higher (*p* < 0.001) with release percentage of 82.51 ± 2.95 %, compared to the Cip release at pH 7.4 (39.10 ± 1.55 %) after 100 h. Over time, the variation in the Cip release percentage at different pH levels significantly escalated, as the Cip release at pH 5.6 at the 100^th^ h was almost double than the Cip release at pH 7.4. The expedited release of Cip at acidic pH was enhanced by the cleavage of the labile succinic acid spacer between Cur and EPS as reported earlier. (11) Moreover, the release profile of free Cip was also determined to validate the sustained Cip release in its loaded form. The release of free Cip exhibited a burst release at pH-7.4 with release of approximately 80 % of Cip within 15 h. This observation indicates that the incorporation of Cip into Cur-EPS micelles enhanced sustained release of Cip. Consequently, encapsulating Cip within micelles can improve its residence time at physiological pH and also boost the Cip release at the biofilm infection site. The kinetics for Cip release from the Cur-EPS micelles at physiological (7.4) and acidic pH (5.6) are further elaborated in the supplementary section (Appendix S6). Additionally, the storage stability of Cip-Cur-EPS is mentioned in the supplementary section (Figure S5).

### 3.4. *In vitro* antibiofilm activity of Cip-Cur-EPS

The minimum biofilm inhibitory concentration (MBIC) and minimum biofilm eradication concentration (MBEC) of free Cip, blank Cur-EPS, and Cur-EPS loaded with Cip against MRSA and *P. aeruginosa* are displayed in the Table 4. It was noted that Cip, in both its loaded and free forms, exhibited a concentration-dependent biofilm inhibitory and dispersion effect for both bacterial strains (Figure S7). These observations can be correlated with the microtiter plate assays depicting the inhibition and eradication of biofilm biomass and viable bacteria within the biofilm formed by both bacterial strains (Figure S6). The blank Cur-EPS micelles demonstrated significantly higher biofilm inhibitory concentrations, which were above 1000 µg/mL for both the bacterial strains. The antibiofilm effect of Cip was enhanced for both bacterial strains when incorporated in Cur-EPS micelles, which was indicated by 4-fold decrease in MBIC and MBEC values compared to free Cip for both the tested bacterial strains. A comparable observation was reported earlier, indicating that Cip, in its encapsulated form, effectively reduced the MBIC by 2- to 4-fold in the examined isolates of *S. aureus* relative to free Cip. (33) Although a high dosage of antibiotics is usually required to mitigate biofilm infections, this can also be correlated with an increased incidence of antibiotic resistance and systemic toxicity. Consequently, incorporating antibiotics such as Cip in Cur-EPS micelles can be advantageous in reducing its dosage while facilitating sustained release at the infection site and boosting the antibiofilm effect of Cip. In addition, Cur-EPS micelles were synthesized to improve solubility and stability and enable pH-responsive release of Cur under acidic conditions. (11) Therefore, the Cip loaded within the Cur-EPS micelles facilitated the release of both Cip and Cur under acidic conditions, which could possibly enhance the antibiofilm effect relative to free Cip. The combination of Cur and Cip has been reported earlier, where a considerable reduction in the minimal inhibitory concentration of Cip was observed. The minimal inhibitory concentration of Cip was reduced by 2-fold for *P. aeruginosa* and 4-fold for MRSA, in comparison to treatment with Cip alone. (34) Additional investigations were conducted to determine the effect of Cip in both its free and loaded forms within Cur-EPS micelles for the eradication of mature biofilms formed by MRSA and *P. aeruginosa*. Further studies were performed using Cip and Cip-Cur-EPS (at the MBIC for Cip-Cur-EPS) to assess improvement in the antibiofilm effect of Cip after being loaded within the Cur-EPS based bioconjugate carrier. The concentration of blank Cur-EPS was kept similar to that of Cip-Cur-EPS, which was used as a control.

**Table 4.** MBIC and MBEC of Cur-EPS, Cip and Cip-Cur-EPS against MRSA and *P. aeruginosa*.

|  | *P. aeruginosa* |  | MRSA |  |
| --- | --- | --- | --- | --- |
| | MBIC<br>($\mu\text{g/mL}$) | MBEC ($\mu\text{g/mL}$)<br>72 h mature biofilm | MBIC<br>($\mu\text{g/mL}$) | MBEC ($\mu\text{g/mL}$)<br>72 h mature biofilm |
| **Cur-EPS** | 1250 | 5000 | 2500 | > 5000 |
| **Cip** | 9.70 | 19.50 | 19.50 | 39.00 |
| **Cip-Cur-EPS** | 2.40 | 4.80 | 4.80 | 9.70 |

### 3.5. Time-kill kinetics for disruption of mature biofilm

In the present study time-kill kinetics experiments were employed to elucidate the time-dependent effects of Cip in both its free and loaded forms on the dispersal of mature biofilms formed by MRSA and *P. aeruginosa* (Figure 5). Cip-loaded Cur-EPS micelles exhibited a time-dependent decrease in the survival of bacterial cells within biofilms formed by both bacterial strains, exhibiting a significant decrease (*p < 0.001*) after treatment for 72 h compared to the control, blank Cur-EPS, and free Cip. Moreover, Cip-Cur-EPS exhibited approximately 1.9- and 3.4 - fold reduction of in log_10_ CFU/mL for *P. aeruginosa* and MRSA, respectively, compared to free Cip after 72 h. Furthermore, from the time-kill kinetics experiments the bactericidal, bacteriostatic, or synergistic activity of the antimicrobial agents can be determined. The reduction in ≥ 3 log_10_ CFU/mL were observed for both the bacterial strains after treatment with Cip-Cur-EPS, compared to the control, which was equivalent to 99.9% killing the viable bacterial cells. (35) Therefore, it was also indicated that the Cip-Cur-EPS exerted bactericidal activity against both the bacterial strains.

**Figure 5.**
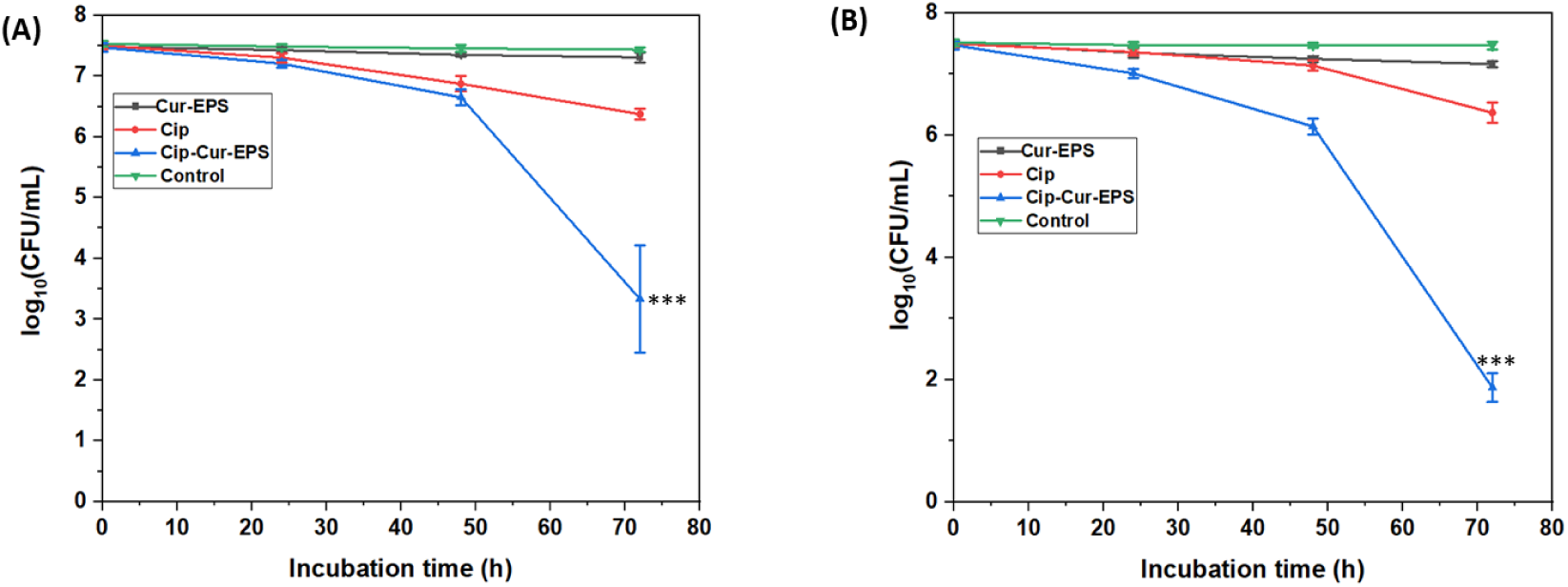
Time-kill kinetics to determine dispersive effects of blank Cur-EPS, free Cip and Cip-Cur-EPS against biofilms formed by (A) MRSA and (B) *P. aeruginosa*. Data is presented as mean values and standard deviation of three independent experiments. Significant levels indicated by asterisks (*\*\*\* p < 0.001*) compared to free Cip at 72 h.

### 3.6. Visualization of biofilm dispersal by CLSM and SEM

An effective strategy for tackling biofilm infections is to disrupt the structure of biofilms in order to uncover cells that can improve the therapeutic effect of antimicrobial drugs. (36) The effect of Cip in both its free and loaded forms on the dispersal of the biofilms formed by MRSA and *P. aeruginosa* was investigated by CLSM and SEM analysis. These microscopy techniques enable visualization and assessment of the 3D structure of biofilm, which offers valuable information regarding the biofilm dispersal post-treatment with test samples. Therefore, this approach provides crucial insights into the effectiveness of the treatment.

The untreated biofilms were distinguished by a uniform cell-matrix distribution over the microscopic field whereas the treatment with Cip-Cur-EPS effectively disrupted the mature biofilms of MRSA and *P. aeruginosa* on glass coverslips compared to blank Cur-EPS micelles and free Cip (Figure 6A, 6B). The anti-biofilm efficacy of free and loaded form of Cip against *P. aeruginosa* and MRSA biofilms was further determined utilizing COMSAT (Figure 6C). The results indicated that in the absence of treatment, the biofilm biovolume were 27.65 ± 1.80 μm³/μm² for MRSA and 21.09 ± 2.15 μm³/μm² for *P. aeruginosa* biofilms, respectively and the average thickness were 34.34 ± 5.27 μm for MRSA biofilms and 30.12 ± 3.56 μm for *P. aeruginosa* biofilms. Following treatment with Cip-Cur-EPS, the thickness and biovolume of the biofilms formed by MRSA and *P. aeruginosa* were significantly reduced (*p < 0.001*) in comparison to the control, blank Cur-EPS micelles and free Cip. The biovolume reduced to 0.54 ± 0.37 μm³/μm² (MRSA) and 0.33 ± 0.17 μm³/μm² (*P. aeruginosa*), whereas the mean thickness reduced to 0.85 ± 0.66 μm (MRSA) and 1.03 ± 0.20 μm (*P. aeruginosa*) after treatment with Cip-Cur-EPS. Furthermore, in comparison to free Cip, the loaded form of Cip within Cur-EPS micelles demonstrated approximately 40- and 65-fold decline in biovolume for MRSA and *P. aeruginosa* biofilms, respectively, and approximately 27- and 19-fold decline in mean thickness for MRSA and *P. aeruginosa* biofilms, respectively.

**Figure 6.**
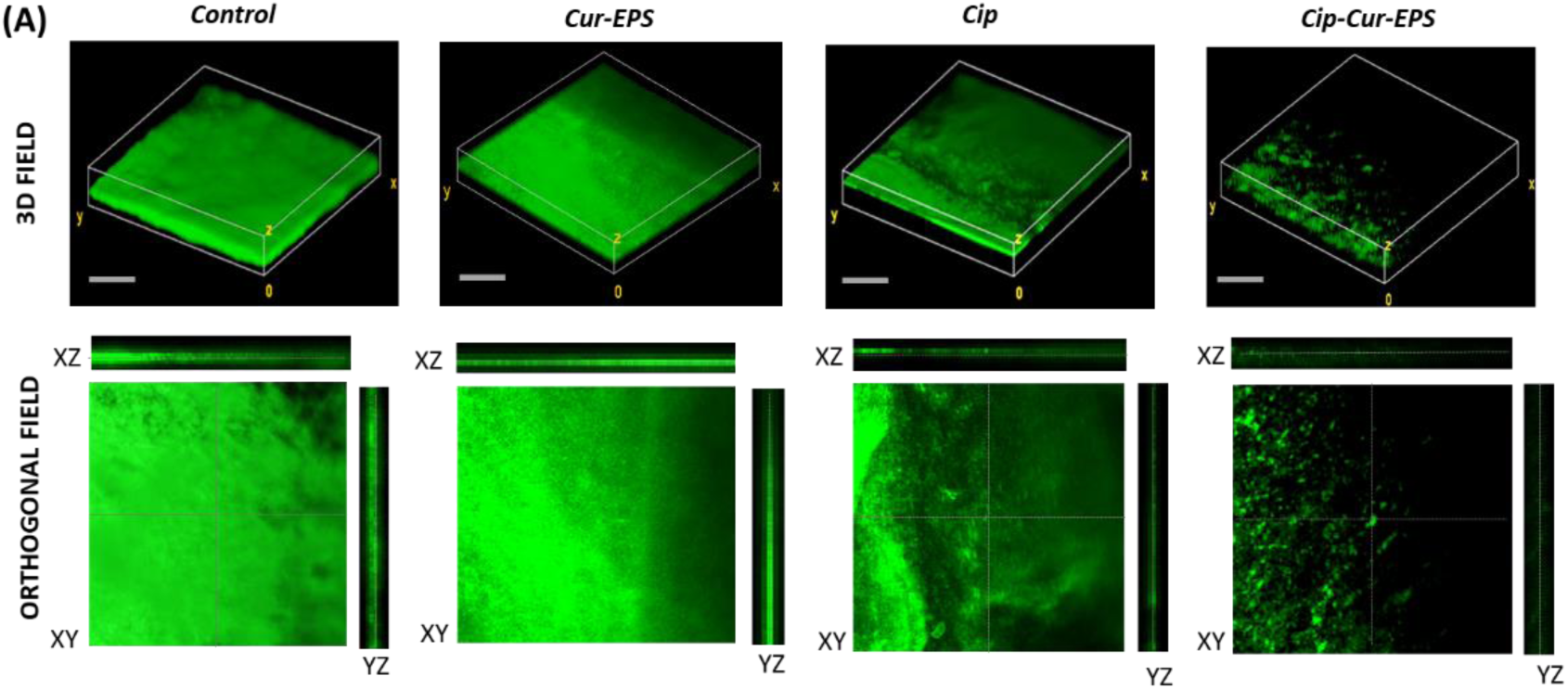
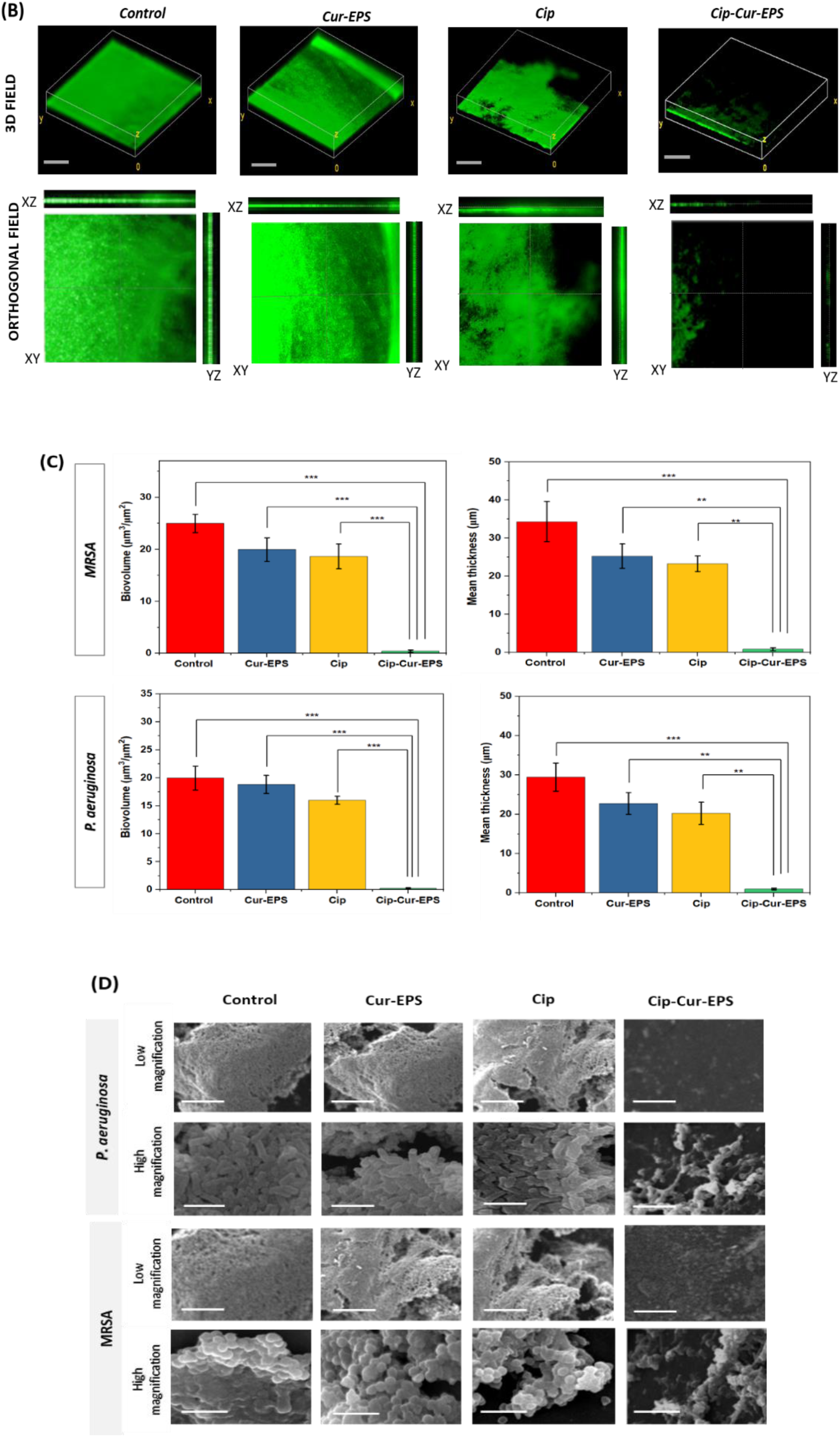
CLSM of biofilm inhibition by (A) MRSA and (B) *P. aeruginosa* treated with blank Cur-EPS, free Cip, and Cip-Cur-EPS. The images illustrate 3D and 2D visualizations at 20 × magnification. Biofilms were stained with acridine orange, causing biofilm biomass appear green when observed with CLSM. Scale bars represent 50 µm. (C) COMSTAT analysis of the MRSA and *P. aeruginosa* biofilms for control and biofilms treated with blank Cur-EPS, free Cip and Cip-Cur-EPS. Data is presented as mean values and standard deviation of three independent experiments. Significant levels indicated by asterisks (*\*\*p < 0.01* and \*\*\**p < 0.001*). (D) SEM image of biofilms formation by *P. aeruginosa* and MRSA at low magnification (450 ×) (scale bar: 50 µm) and high magnification (6000 ×) (scale bar: 3 µm).

Furthermore, the effects of Cip in its free and loaded form on biofilm structures of MRSA and *P. aeruginosa* were also visualized by SEM (Figure 6D). At low magnification, the control group cells merged to form macro-colonies, demonstrating a mature biofilm with a complex architecture. After 72 h treatment with Cip-Cur-EPS, the biofilm architecture was significantly compromised. At elevated magnification, all cells in the control group remained unaltered. Rod-shaped (*P. aeruginosa*) and spherical-shaped (MRSA) cellular aggregates were observed in the control, blank micelle, and free Cip, showing that the cells within the biofilm were thriving. Following treatment with Cip-Cur-EPS for 72 h, distorted cells were observed in case of both the bacterial strains. Consequently, these outcomes indicate Cip contained within Cur-EPS micelles exhibited considerably higher ability in eradicating biofilm biomass compared to its free form.

### 3.7. *In vitro* biofilm dispersal effects of on the urinary catheter pieces under static condition

The formation of biofilm on urinary catheters plays a vital role in the progression of catheter-associated urinary tract infections (CAUTI). Repeated insertion of indwelling catheters has increased the incidence of CAUTI, which are influenced by various factors, such as the specific pathogens present in the hospital, immune function, microbiota. and infection history of the patient. (37) Patients suffering from CAUTI frequently receive treatments with high dosage of antibiotics, urinary catheter replacement, and routine bladder irrigation to promote bacterial eradication. However, such approaches often cause pain and increased suffering in patients, while also promoting the development of drug-resistant bacteria and antibiotic-related side effects, consequently complicating treatment and racking up the expenses. (37) Therefore, an alternative approach is required that can inhibit the proliferation and colonization of pathogenic bacteria, and reduce side effects pertaining to high doses of antibiotic. (38) In this study, the effectiveness of Cip-loaded Cur-EPS micelles in eradicating mature biofilms of MRSA and *P. aeruginosa* was evaluated under *in vitro* conditions on urinary catheter surfaces in the presence of AUM.

Initially the bacterial growth and biofilm formation was monitored in AUM which closely resembles to the human urine. Figure S8 indicates that the biofilm formation of MRSA and *P. aeruginosa* in AUM were comparable to that observed in TSB. The biofilm biomass formed by MRSA and *P. aeruginosa* was evaluated on treated and untreated catheters under static conditions with crystal violet assay. Figure 8A presents the CV-stained biofilm biomass on the silicon catheter fragments. The decrease in staining was regarded as an indicator of diminished biofilm formation on the catheter surface, which was observed after treatment with Cip-Cur-EPS. The eradication percentages of biofilm biomass were subsequently assessed for the treated catheter segments from the absorbance of crystal violet stain at 570 nm. The catheters treated with Cip-Cur-EPS significantly (*p < 0.001*) disrupted the biofilms formed by MRSA and *P. aeruginosa* compared to the control, blank Cur-EPS, and free Cip after the treatment period for 72 h (Figure 8B). The Cip-Cur-EPS eradicated biofilms formed by MRSA and *P. aeruginosa* by 82.04 ± 3.13 % and 90.56 ± 2.36 % respectively. Treatment with Cip-Cur-EPS increased the biofilm eradication rates by approximately 4.5- and 5.2-fold for MRSA and *P. aeruginosa* respectively, in comparison to free Cip. Additionally, the impact of treatments on viable cells within the biofilms produced by MRSA and *P. aeruginosa* were assessed by determining the log reduction values. The viability of both tested bacterial strains deteriorated following treatment with Cip-Cur-EPS, as evidenced by a significant (*p < 0.001*) increase in log reduction values compared to the control, blank Cur-EPS, and free Cip (Figure 8C). The treatment with Cip-Cur-EPS exhibited considerably high LRV of 1.63 ± 0.15 and 2.21 ± 0.42 for MRSA and *P. aeruginosa* respectively. Moreover, the log reduction values for MRSA and *P. aeruginosa* increased by approximately 10- and 16-fold respectively after treatment with Cip-Cur-EPS relative to free Cip.

**Figure 8.**
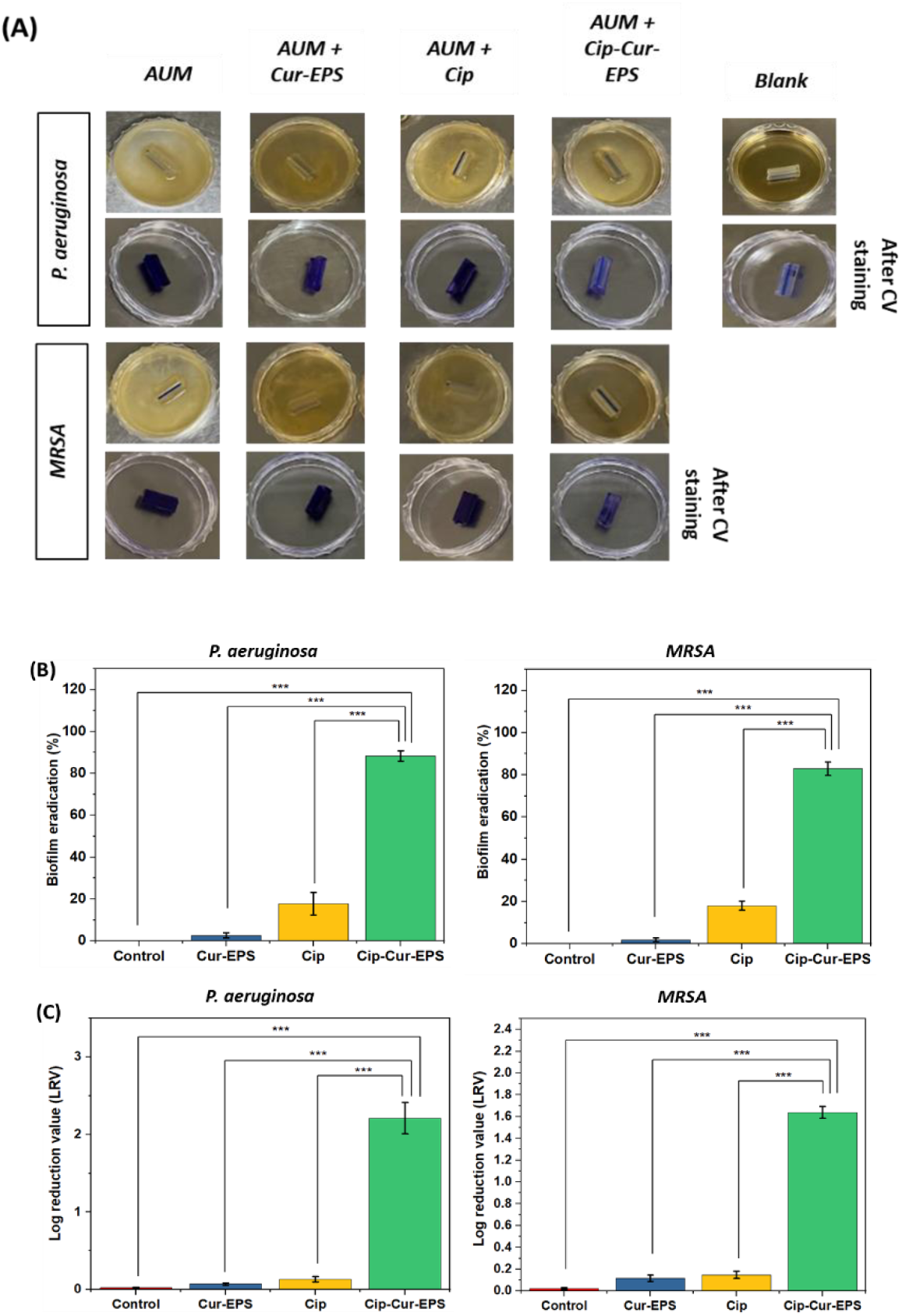
(A) The biofilm formation *P. aeruginosa* and MRSA under static conditions on untreated and treated catheter pieces determined by CV staining. (B) Biofilm biomass eradication percentage of blank Cur-EPS, free Cip, and Cip-Cur-EPS against *P. aeruginosa* and MRSA. (C) Log reduction values after treatment with blank Cur-EPS, free Cip, and Cip-Cur-EPS against P. aeruginosa and MRSA. Statistically significant difference (p < 0.001 are denoted by ***), values are presented as mean ± SD, n = 3.

The aforementioned results were further corroborated by microscopic examination of the biofilm utilizing SEM (Figure 9). A dense aggregation of intact and closely packed cells appeared on the surface of the untreated silicone catheters. Treatment of mature biofilms on the catheter surface with Cip in its loaded form caused a significant damage to the biofilms compared to free Cip demonstrating superior effectiveness of Cip-Cur-EPS in disrupting the structural integrity and reducing cell aggregates of MRSA and *P. aeruginosa*. Moreover, effective biofilm treatment includes both biofilm dispersion and the prevention of bacterial recolonization on the surface post-treatment. Treatment of the biofilms with Cip-Cur-EPS was also capable of inhibiting the re-growth of *P. aeruginosa* and MRSA cells on the catheter surface, in contrast to free Cip (Figure 10).

**Figure 9.**
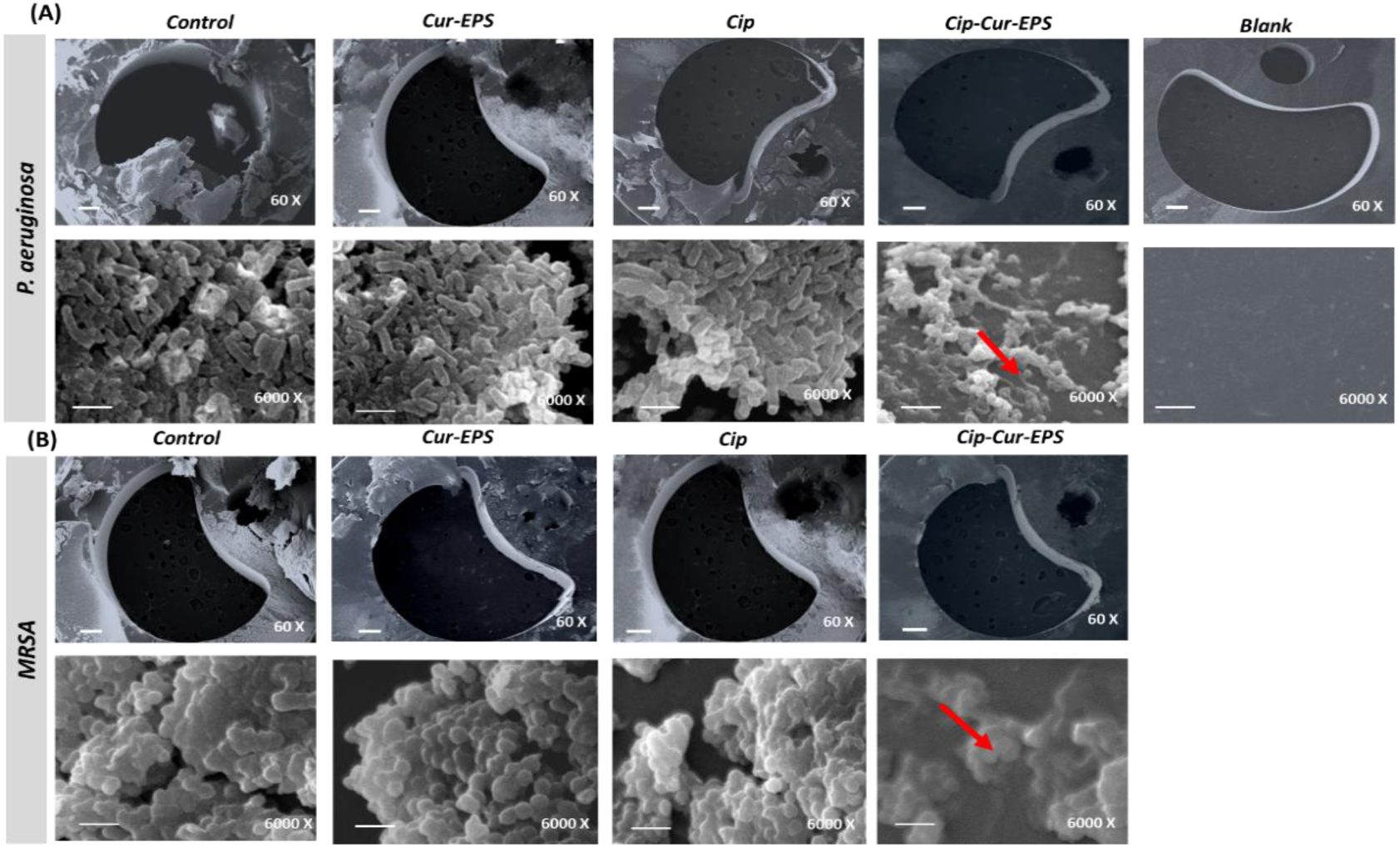
Visualization of biofilm thickness by SEM on catheter cross sections after treatment with blank micelle, free Cip and Cip-Cur-EPS at low magnification (scale bar: 100 µm) and high magnification (scale bar: 2 µm).

**Figure 10.**
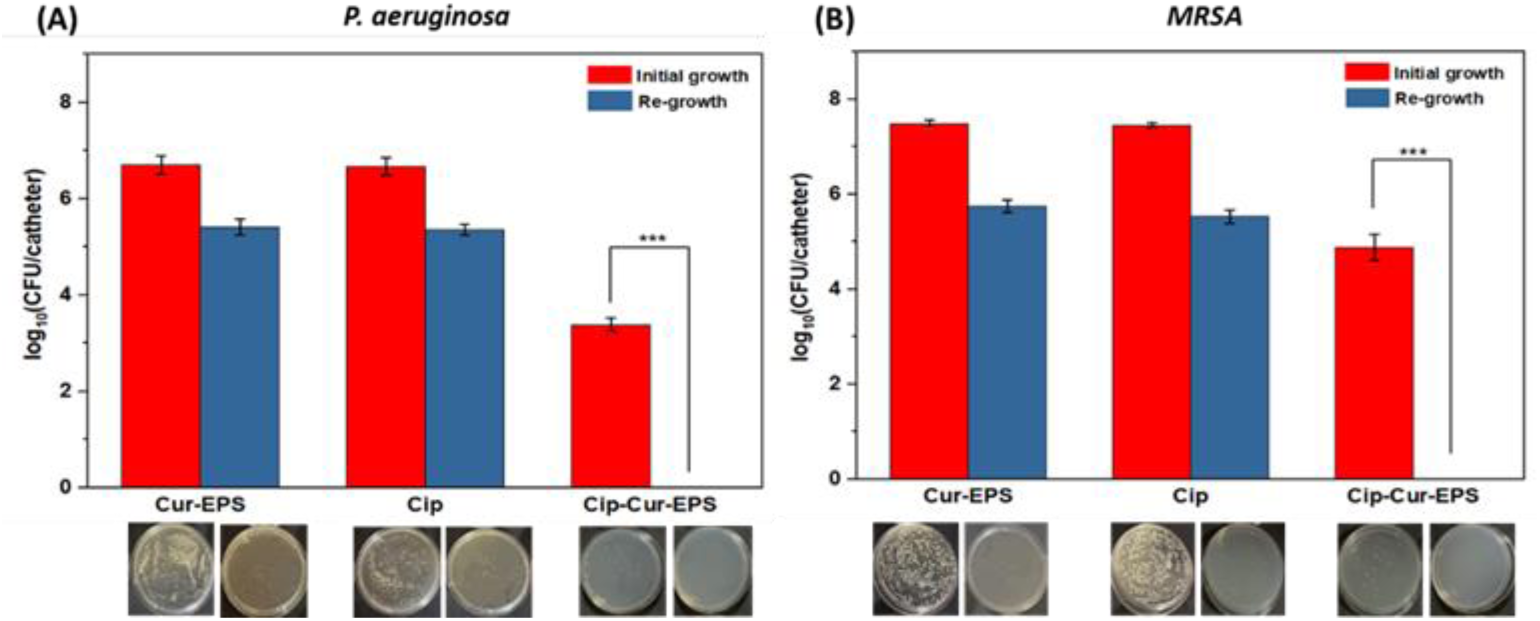
Effect of treatments on regrowth capability of *P. aeruginosa* (A) and MRSA (B). For each strain, data are expressed as log (CFU/catheter). Representative agar plate images are presented below the graph. Statistically significant difference is denoted by *** (*p <* 0.001) and the values are represented as mean ± SD, n = 3.

### 3.8. Biofilm dispersal tests under dynamic conditions in a model of catheterized human bladder

Although the biofilm dispersal effect of Cip-Cur-EPS MRSA and *P. aeruginosa* were assessed under static conditions, it does not accurately reflect the actual circumstances encountered during catheterization. The development of experimental models that simulate the urodynamic settings of human bladder is essential for investigating the pathogenic mechanisms of catheter-associated bacterial biofilm infections. Consequently, the biofilm dispersal effects of Cip-Cur-EPS under dynamic settings were further evaluated in a model of catheterized human bladder (Figure 11A). A noteworthy decrease in the biofilm biomass of MRSA and *P. aeruginosa* were observed after 72 h, which was indicated by significant increase (*p <* 0.001) in the biofilm eradication percentage after treatment with Cip-Cur-EPS compared to control, blank Cur-EPS micelles and free Cip (Figure 11B). Under hydrodynamic conditions Cip-Cur-EPS eradicated the biofilms formed by MRSA and *P. aeruginosa* with biofilm eradication of 79.53 ± 3.58 % and 85.94 ± 4.98 % respectively. These observations were almost consistent with the static experiments. The treatment with Cip-Cur-EPS led to 7.9- and 9.4-fold increases in the percentages of biofilm removal by MRSA and P. aeruginosa, respectively, compared to free Cip. The qualitative results that were depicted by the photographs of urinary catheters aligned with the quantitative data (Figure 11C). Consequently, these results indicate that Cip-loaded Cur-EPS micelles facilitated dispersal of mature biofilm within a simulated bladder environment, which has significant implications for urinary tract infection management. The ability of these Cip-loaded Cur-EPS micelles to effectively disrupt established biofilms suggests a promising approach for combating persistent infections associated with medical devices such as urinary catheters. This enhanced biofilm removal capability could potentially lead to more effective treatments, reduced infection rates, and improved patient outcomes in clinical settings where biofilm-related infections pose a significant challenge.

**Figure 11.**
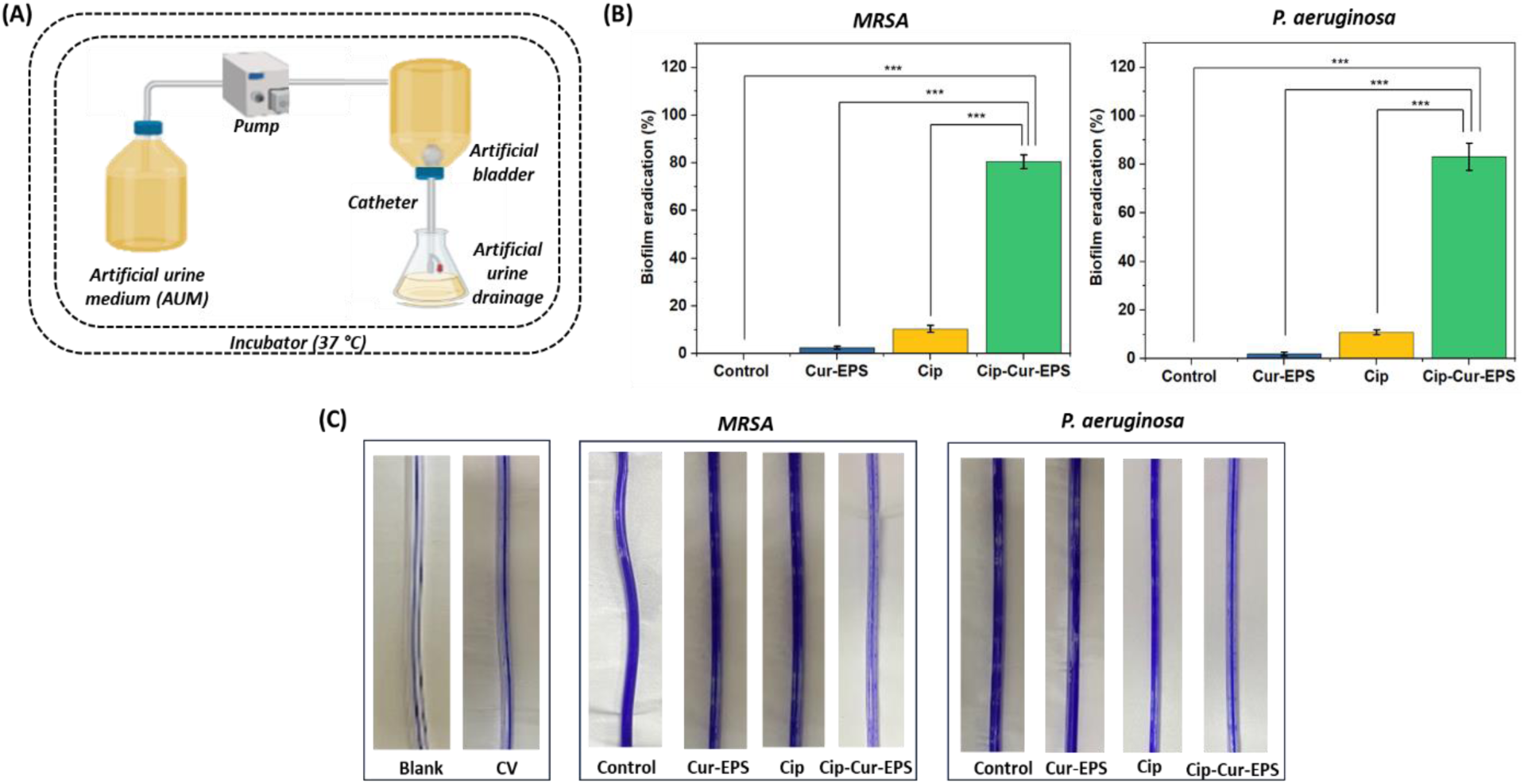
(A) Schematics representation of the *in vitro* model of catheterized bladder. (B) Bacterial biofilms visual image on catheter surface by crystal violet staining. (C) Biofilm eradication percentage against mature biofilms formed by *P. aeruginosa* and MRSA urinary catheters by blank Cur-EPS, free Cip, and Cip-Cur-EPS after 3 days. Statistically significant difference (*p < 0.001* are denoted by ***), values are presented as mean ± SD, n = 3.

## Conclusion

In this study, optimized formulation of Cip-loaded Cur-EPS micelles were prepared, which improved the solubility of Cip at physiological pH. The Cip-Cur-EPS micelles demonstrated enhanced antibiofilm effects relative to free Cip, by virtue of their pH-responsive delivery of Cip, facilitating their utilization in pathological regions such as biofilm infection sites. The results confirmed that Cip in its encapsulated form within Cur-EPS micelles surpassed the antibiofilm effect of free Cip in reducing overall biofilm biomass, for both examined bacterial strains. Furthermore, the effects of Cip-Cur-EPS micelles in mitigating biofilm formation on the surface of medical devices, such as urinary catheters, were assessed. Cip-loaded Cur-EPS micelles were considerably more effective than free Cip in eradicating biofilms formed on the surface of a urinary catheter by leveraging the low pH within the biofilms. The findings from this study have laid a foundation for application of Cur-EPS micellar bioconjugates as carriers for antibiotics that can potentially improve therapeutic effect of antibiotics by efficiently eradicating biofilm related infections in patients with chronic urinary tract infections. Although in this work, the biofilm dispersive effect of Cip-Cur-EPS have been studied, further extensive studies are required to uncover the bactericidal mechanism of this formulation and their effectiveness for polymicrobial infections in *in vitro* and *in vivo* models.

## AUTHOR INFORMATION

### Authors

**Chandrika Gupta -** *Department of Biotechnology, Indian Institute of Technology Kharagpur, West Bengal, India;*

**Sudip Kumar Ghosh** *- Department of Biotechnology, Indian Institute of Technology Kharagpur, West Bengal, India;*

### Author contributions

Writing - original draft, methodology, validation: C.G.; Validation, Supervision: S.K.G. conceptualization, writing - review & editing, validation, resources, supervision: R.S.

## Acknowledgments

The experimental facilities and resources for this study was supported by the Indian Institute of Technology Kharagpur. CG is grateful to the Ministry of Human Resources and Development, Government of India, for the Doctoral Fellowship.

