## Supplementary information for "Performance Evaluation of Ciprofloxacin loaded Micellar Bioconjugate Carrier for Eradication of Bacterial Biofilms"

**Appendix S1:**

**Release kinetics models**

The equations for the kinetic models to determine the Cip release profile as reported earlier, are mentioned as follows: (1)

Zero order model:

$$C_t = C_0 + k_0 t \quad (1)$$

$Ct$ is the amount of drug released at time $t$, $C_0$ is the initial concentration of drug at time t = 0, $k_0$ is the zero-order rate constant. This model represents drug-release data by estimating the cumulative amount of Cip released over a period of time.

First order model:

$$log\ (C_0) – log(C_t) = k_1 t\ /2.303 \quad (2)$$

$k_1$ is the first-order release constant, $t$ is the time of drug release. Here, the change in drug concentration is a function of the instantaneous concentration.

Higuchi model:

$$M_t\ /\ M_\infty = K_h t^{1/2} \quad (3)$$

$M_t / M_\infty$ is a fraction of drug released at time $t$, $M_t$ represents the amount of drug released over time, $M_\infty$ represents the amount of drug at equilibrium and $K_h$ is the Higuchi release kinetic constant. Drug release in this context is described as a diffusion process that follows Fick's law and is reliant on the square root of time.

Hixson-Crowell model:

$$C_0^{1/3} - Ct^{1/3} = k_{HC}t \quad (4)$$

where $k_{HC}$ is the Hixon-Crowell dissolving rate constant. This model depicts a system where the cube root of the amount of drug released is directly proportional to time. This system specifically involves changes in its surface over time.

Korsmeyer-Peppas model:

$$M_t / M_\infty = Kkpt^n \quad (5)$$

$n$ is the release exponent indicating the Cip release mechanism, $K_{kp}$ is the Korsmeyer-Peppas release rate constant and denotes the geometric and structural features of the drug and $n$ is the release exponent indicating the drug release mechanism. This model is also known as the "power law", and it is a diffusion-based semi-empirical approach widely used to demonstrate the main transport processes involved in the release, either through diffusion or swelling.

Peppas-Sahlin model:

$$M_t / M_\infty = k_1 t^m + k_2 t^{2m} \quad (6)$$

$k_1$ is the Fickian kinetic constant for Cip release, $k_2$ is the erosion rate constant (kinetic constant for Case-II transport), and $m$ is the diffusional exponent for systems of any geometrical shapes displaying drug-controlled release. This model is employed to analyse both Fickian and non-Fickian transfer from the delivery systems.

**Appendix S2:**

**S2.1. Antibiofilm activity of Cur-EPS, Cip and Cip-Cur-EPS**

The minimal biofilm inhibition concentration (MBIC) and minimal biofilm eradication concentration (MBEC) for Cur-EPS micelle, Cip and Cip-Cur-EPS were determined by the protocol described by Haney et al. with minor modifications. (2) For this study, 90 µL of bacterial cultures ($10^6$ CFU/mL) prepared in Tryptic Soy Broth (TSB) was inoculated into sterile flat-bottomed 96-well microtiter plates with aqueous aliquots of 10 µL of Cip, Cip-Cur-EPS (with equivalent amount of Cip at concentration range of 0.6 – 39 µg/mL) and Cur-EPS (19.5 – 5000.0 µg/mL) respectively, into each well and was incubated for 72 h at 37 °C under static condition. The wells containing diluted bacterial suspension without the test samples were designated as growth controls. After the incubation period the cultures were removed, and the wells were washed with PBS to remove any non-adherent cells. The inhibition of biofilm in the wells after 72 h was assessed in terms of the biofilm biomass and metabolic activity of bacteria embedded within the biofilm. Therefore, the experiments were simultaneously performed in two different microtiter plates. The concentration dependent inhibition of biofilm biomass was determined by crystal violet assay. Following the incubation period, the broth containing planktonic cells were removed and wells were rinsed three times with PBS. The adherent biofilms were heat fixed at 60 °C for 45 min, and then stained with 100 µL of 1% crystal violet for 5 min. The unbound stain was removed and the wells were rinsed thrice with PBS and the remaining dye was dissolved in 95% ethanol for 15 min. The absorbance was acquired at a wavelength of 540 nm using a spectrophotometer with a plate reader (Thermoscientific Multiskan Go, USA). Furthermore, the inhibition of bacterial metabolic activity within the biofilm was determined by TTC reduction assay. After the incubation period, the culture was removed, and the microtiter wells were washed with PBS to remove planktonic cells. 20 µL of the TTC reagent, with a concentration of 0.5 mg/mL, was added to the microtiter plates and were placed in a dark environment at 37 °C for a duration of 30 min. Following the incubation period, the TTC was substituted with 100 µL of DMSO in order to dissolve the formazan complex. The optical density of each well was quantified at a wavelength of 490 nm using a microplate reader. MBIC was considered as the lowest amount of test sample that completely prevents biofilm formation visually and by measuring the absorbance at 490 nm. Further, different concentrations of samples, were introduced to each well containing a 72-h old biofilm. Subsequently, the plates were placed in an incubator at 37 °C for 24 h. MBEC in terms of eradication of biomass and viable cells were assessed using the crystal violet assay and TTC assay as mentioned previously. The MBEC was determined as the lowest concentration of antimicrobial agents needed to achieve complete eradication of preformed biofilm. The results were also quantified as percentages of biofilm inhibition and eradication relative to the control.

**S2.2. Time-kill study of bacterial biofilm disruption of mature biofilm**

To determine the time-kill study for disruption of bacterial biofilm, a bacterial suspension of $10^6$ CFU/mL from an overnight culture was added to wells of a microplate containing TSB. The plates were incubated at 37 °C for 72 h to facilitate the development of biofilm. After incubation, the media containing planktonic cells were removed from the wells, and the biofilm was rinsed with PBS. Following treatment, blank micelle, Cip and Cip-Cur-EPS (at equivalent concentration of Cip) and incubation at three specific time intervals: 24, 48 and, 72 h at a 37 °C the biofilm was further scraped and the viable bacterial cells were determined as mentioned in the previous section. Untreated wells containing only the inoculated bacterial culture were considered as growth controls.

**Appendix S3:**

**SEM of bacterial biofilms**

The impact of blank Cur-EPS, free Cip, and Cip-loaded Cur-EPS on biofilm matrix structure was examined utilizing a scanning electron microscope (Quanta 200 SEM, FEI, Hillsboro,

Oregon, USA). Bacterial biofilms were cultivated on glass cover slips in a 24-well microplate for 48 hours at 37 °C. Subsequently, biofilms were dried using a graduated series of ethanol concentrations and analysed via SEM.

**Appendix S4:**

**S4.1. Preparation of Artificial urine medium**

The artificial urine medium (AUM) was prepared according to the formulation that was reported earlier. (3) The AUM composition included: urea 13 g/L, uric acid 0.33 g/L, creatinine 1.2 g/L, sodium chloride 2.9 g/L, ammonium chloride 3.0 g/L, sodium sulphite 3.0 g/L, disodium hydrogen orthophosphate 2.5 g/L and potassium dihydrogen orthophosphate 2.5 g/L. The pH of AUM was adjusted to 6.5 with 1 M NaOH or HCl and was further sterilized by autoclaving at 121°C, 15 psi for 15 min. All the chemicals were procured from SRL Pvt. Ltd. Mumbai, India.

**S4.2. Bacterial growth and biofilm formation in Artificial urine medium (AUM)**

*P. aeruginosa* and *MRSA* were preliminary tested for their growth in AUM. Initially, bacterial strains were inoculated in TSB and incubated at 37 °C, for 24 h. After the incubation period the bacterial suspensions were centrifuged (4000 rpm, 10 min) and were resuspended in PBS for subsequent experiments. Aliquots of the diluted suspensions (~$10^6$ CFU/mL) were pipetted into the wells of a 24-well microtiter plate containing 1 mL of AUM and the plate was incubated for 24 h with gentle shaking at 37 °C. Afterwards, the culture in each well was diluted and 50 μL was inoculated on MHA plates to determine the colony forming units (CFU/mL). To access the biofilm formation in AUM, bacterial cultures (~$10^6$ CFU/mL) were added into wells of a 24-well plate containing AUM. The plates were incubated at 37 °C. After incubation period of 72 h, the wells were washed in PBS to remove planktonic bacterial cells. Subsequently, PBS was added to each well to dislodge the biofilm by pipetting followed by further ultrasonication (46 kHz, 2 min). In order to determine the quantity of live bacteria, the sonicated suspensions were diluted in PBS and spread on TSA plates. Following a 24 h incubation period at a 37 °C, the CFU/mL was determined. Biomass analysis was carried out using CV staining as mentioned in the previous section.

**Table S1:** The experimental runs and corresponding responses in terms of the particle size, PDI and EE.

| | **Independent variables (Factors)** | | | **Dependant variables (Responses)** | | | | | |
|---|---|---|---|---|---|---|---|---|---|
| | A: Cur-EPS concentration (mg/mL) | B: Tween 80 (%) | C: Water:DMSO | X1: Particle size (nm) | | X2: PDI | | X3: EE (%) | |
| | | | | PV | EV | PV | EV | PV | EV |
| 1 | 0 | 0 | -1.68 | 316.95 | 315.23 ± 0.23 | 0.55 | 0.54 ± 1.42 | 47.12 | 45.92 ± 2.43 |
| 2 | 0 | 0 | 1.68 | 304.34 | 303.17 ± 1.45 | 0.40 | 0.41± 1.13 | 86.97 | 85.34 ± 2.56 |
| 3 | 1 | 1 | 1 | 346.10 | 345.23 ± 0.63 | 0.39 | 0.40 ± 0.45 | 76.50 | 78.45 ± 1.75 |
| 4 | 0 | 0 | 0 | 325.74 | 327.42 ± 1.25 | 0.41 | 0.40 ± 0.34 | 78.44 | 77.09 ± 1.65 |
| 5 | 0 | 1.68 | 0 | 326.19 | 325.03 ± 0.78 | 0.43 | 0.42 ± 0.56 | 63.91 | 63.87± 2.35 |
| 6 | 1 | -1 | -1 | 369.46 | 369.56 ± 1.35 | 0.62 | 0.63 ± 0.34 | 63.17 | 64.45 ± 1.43 |
| 7 | -1 | 1 | 1 | 305.21 | 308.00 ± 1.32 | 0.38 | 0.37 ± 0.76 | 68.35 | 69.00 ± 2.06 |

| | | | | | | | | | |
|---|---|---|---|---|---|---|---|---|---|
| 8 | 0 | 0 | 0 | 325.74 | 325.23 ± 1.65 | 0.41 | 0.43 ± 0.32 | 78.44 | 79.03 ± 1.54 |
| 9 | 1 | -1 | 1 | 357.97 | 358.56 ± 1.45 | 0.53 | 0.53 ± 0.65 | 80.14 | 80.45 ± 2.78 |
| 10 | 0 | -1.68 | 0 | 341.10 | 339.10 ± 0.89 | 0.69 | 0.71 ± 1.12 | 62.18 | 60.34 ± 2.56 |
| 11 | 1.68 | 0 | 0 | 392.91 | 395.32 ± 1.75 | 0.51 | 0.49 ± 1.85 | 68.98 | 69.67 ± 2.45 |
| 12 | 1 | 1 | -1 | 353.09 | 353.00 ± 1.98 | 0.53 | 0.54 ± 0.45 | 63.03 | 62.21 ± 1.87 |
| 13 | 0 | 0 | 0 | 325.74 | 325.43 ± 2.67 | 0.41 | 0.42 ± 1.56 | 78.44 | 79.04 ± 3.12 |
| 14 | -1 | 1 | -1 | 308.71 | 311.04 ± 2.65 | 0.47 | 0.47 ± 0.78 | 53.38 | 55.32 ± 2.84 |
| 15 | -1 | -1 | 1 | 306.58 | 309.11 ± 2.12 | 0.60 | 0.58 ± 1.09 | 74.49 | 77.34 ± 1.74 |
| 16 | 0 | 0 | 0 | 325.74 | 325.54 ± 1.75 | 0.41 | 0.42 ± 1.25 | 78.44 | 78.56 ± 1.32 |
| 17 | -1 | -1 | -1 | 314.57 | 318.00 ± 1.52 | 0.65 | 0.63 ± 1.05 | 56.02 | 56.09 ± 2.45 |
| 18 | 0 | 0 | 0 | 325.74 | 327.33 ± 1.75 | 0.41 | 0.40 ± 0.43 | 78.44 | 78.00 ± 1.76 |
| 19 | 0 | 0 | 0 | 325.74 | 326.15 ± 0.56 | 0.41 | 0.41± 0.72 | 78.44 | 80.23 ± 0.65 |

**Table S2:** ANOVA statistical analysis for full factorial model for the particle size.

| Source | Degree of Freedom | Sum of squares | Mean square | F-value | p-value |
|---|---|---|---|---|---|
| **Model** | 9 | 10325.30 | 1147.26 | 140.49 | **0** |
| **Linear** | 3 | 8289.80 | 2763.27 | 338.39 | **0** |
| **A** | 1 | 7829.60 | 7829.59 | 958.82 | **0** |
| **B** | 1 | 268.40 | 268.42 | 32.87 | **0** |
| **C** | 1 | 191.80 | 191.81 | 23.49 | **0.001** |
| **Square** | 3 | 1964.20 | 654.72 | 80.18 | **0** |

| | | | | | |
|---|---|---|---|---|---|
| **$A^2$** | 1 | 1304.20 | 1304.16 | 159.71 | **0** |
| **$B^2$** | 1 | 112.60 | 112.61 | 13.79 | **0.004** |
| **$C^2$** | 1 | 410.40 | 410.38 | 50.26 | **0** |
| **Two-way interaction** | 3 | 71.40 | 23.79 | 2.91 | **0.087** |
| **AB** | 1 | 55.10 | 55.12 | 6.75 | **0.027** |
| **AC** | 1 | 6.10 | 6.12 | 0.75 | **0.407** |
| **BC** | 1 | 10.10 | 10.12 | 1.24 | **0.292** |
| **Error** | 10 | 81.70 | 8.17 | | |
| **Lack of fit** | 5 | 76.80 | 15.37 | 15.89 | **0.064** |
| **Pure error** | 5 | 4.80 | 0.97 | | |
| **Total** | 19 | 10407 | | | |
| **$R^2$** | 0.99 | **Adj. $R^2$** | 0.98 | **Pred. $R^2$** | 0.95 |

**Table S3:** ANOVA statistical analysis for full factorial model for the PDI.

| Source | Degree of Freedom | Sum of squares | Mean square | F-value | p-value |
|---|---|---|---|---|---|
| **Model** | 9 | 0.172263 | 0.01914 | 51.56 | **0** |
| **Linear** | 3 | 0.112362 | 0.037454 | 100.9 | **0** |
| **A** | 1 | 0.00019 | 0.00019 | 0.51 | **0.491** |
| **B** | 1 | 0.085047 | 0.085047 | 229.11 | **0** |
| **C** | 1 | 0.027124 | 0.027124 | 73.07 | **0** |
| **Square** | 3 | 0.055064 | 0.018355 | 49.45 | **0** |
| **$A^2$** | 1 | 0.018763 | 0.018763 | 50.55 | **0** |

| | | | | | |
|---|---|---|---|---|---|
| **$B^2$** | 1 | 0.038958 | 0.038958 | 104.95 | **0** |
| **$C^2$** | 1 | 0.005865 | 0.005865 | 15.8 | **0.003** |
| **Two-way interaction** | 3 | 0.004837 | 0.001613 | 4.34 | **0.033** |
| **AB** | 1 | 0.002813 | 0.002813 | 7.58 | **0.02** |
| **AC** | 1 | 0.001012 | 0.001012 | 2.73 | **0.13** |
| **BC** | 1 | 0.001012 | 0.001012 | 2.73 | **0.13** |
| **Error** | 10 | 0.003712 | 0.000371 | | |
| **Lack of fit** | 5 | 0.002979 | 0.000596 | 4.06 | **0.075** |
| **Pure error** | 5 | 0.000733 | 0.000147 | | |
| **Total** | 19 | 0.175975 | | | |
| **$R^2$** | 0.97 | **Adj. $R^2$** | 0.96 | **Pred. $R^2$** | **0.94** |

**Table S4:** ANOVA statistical analysis for full factorial model for the EE.

| Source | Degree of Freedom | Sum of squares | Mean square | F-value | p-value |
|---|---|---|---|---|---|
| **Model** | 9 | 1773.46 | 197.051 | 66.81 | **0** |
| **Linear** | 3 | 1103.97 | 367.99 | 124.77 | **0** |
| **A** | 1 | 199.73 | 199.727 | 67.72 | **0** |
| **B** | 1 | 33.56 | 33.561 | 11.38 | **0.007** |
| **C** | 1 | 870.68 | 870.682 | 295.2 | **0** |
| **Square** | 3 | 659.11 | 219.703 | 74.49 | **0** |
| **$A^2$** | 1 | 455.07 | 455.074 | 154.29 | **0** |
| **$B^2$** | 1 | 254.84 | 254.843 | 86.4 | **0** |

| | | | | | |
|---|---|---|---|---|---|
| **$C^2$** | 1 | 43.15 | 43.147 | 14.63 | **0.003** |
| **Two-way interaction** | 3 | 10.37 | 3.458 | 1.17 | **0.368** |
| **AB** | 1 | 3.12 | 3.125 | 1.06 | **0.328** |
| **AC** | 1 | 1.12 | 1.125 | 0.38 | **0.551** |
| **BC** | 1 | 6.12 | 6.125 | 2.08 | **0.18** |
| **Error** | 10 | 29.49 | 2.949 | | |
| **Lack of fit** | 5 | 23.99 | 4.799 | 4.36 | **0.066** |
| **Pure error** | 5 | 5.5 | 1.1 | | |
| **Total** | 19 | 1802.95 | | | |
| **$R^2$** | 0.98 | **Adj. $R^2$** | 0.97 | **Pred. $R^2$** | 0.94 |

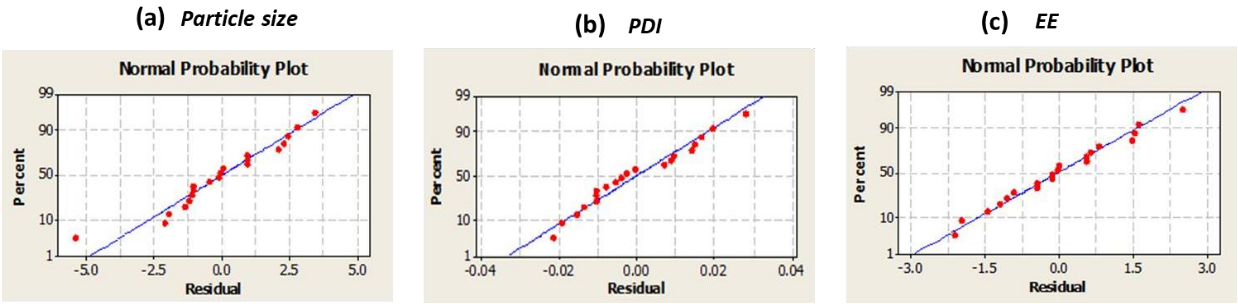

**Figure S1.** Normal probability plot for residuals for (a) particle size, (b) PDI and (c) EE.

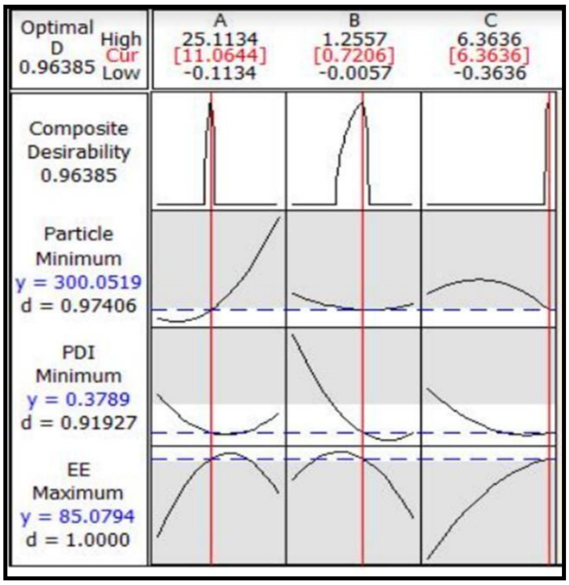

**Figure S2.** Process optimization curve for obtaining the target value.

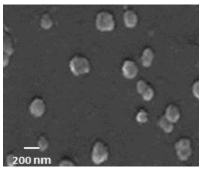

**Fig.S3.** FESEM image of Cip-Cur-EPS

**Appendix S5:**

Mathematical models were applied to describe the release of Cip from the Cur-EPS micelles at physiological and acidic pH. The mechanism of release profiles was analyzed by using several recognized kinetic models, including zero-order, first-order, Higuchi, Hixson-Crowell, Korsmeyer-Peppas and Peppas-Sahlin models. The released fraction was plotted against time (t) to comprehend the release behavior for the Cip release at pH 7.4 and pH 5.6. The diverse parameters of different release models along with the $R^2$ (regression coefficient) and the AIC (Akaike information criterion) values were presented in Table S5. The AIC is a useful metric for comparing models fitted to the same dataset. The model with the lowest AIC is considered as the most appropriate fit for the data. (4) The maximum $R^2$ values (the best linearity) and lowest AIC was found in Peppas-Sahlin model followed by Korsmeyer-Peppas model. The fit of Peppas-Sahlin and Korsmeyer-Peppas models to the experimental release data for Cip from Cur-EPS micelles at pH 7.4 and 5.6 are depicted in Figure S4. The constants $k_1$, $k_2$ and diffusion exponent m were calculated from the Peppas-Sahlin model. The $k_1$ and $k_2$ kinetic constants indicate the Case I transport (Fickian diffusion) and Case II transport (dissolution and relaxation of polymer chains) respectively. When $k_1 > k_2$, it indicates that the drug release is facilitated by Fickian diffusion (Case I diffusion). Conversely, if $k_2 > k_1$, it signifies the dominance of Case II relaxation. When $k_1 = k_2$, the release follows both diffusion and erosion. (1) The results depicted in Table S5 showed that in the present study, the kinetic constant $k_1$ for Fickian contribution was higher than the kinetic constant $k_2$ denoting Case II contribution. This observation reveals that Cip release from Cur-EPS micelles was governed by Fickian diffusion at both pH 7.4 and 5.6. Also, the negative values for $k_2$ suggests that the erosion rate constant does not play a substantial role in the Cip release. Moreover, the experimental release data for Cip also demonstrated a significant correlation and lower AIC values with the Korsmeyer-Peppas model at both pH. The diffusion exponent (n) was calculated for the release of Cip from the micelle. The values obtained were 0.37 and 0.50 at pH 7.4 and 5.6, respectively, using the Korsmeyer-Peppas model. Considering that the values of $n \leq 0.5$, the measured diffusion exponent values indicate Fickian diffusion. Similar findings were reported earlier for the release profile of Cip through hydrophilic matrices. (5)

**Table 4.8:** Results of fitting the release profiles with different kinetic models.

| **Model** | | **Physiological medium (pH 7.4)** | **Acidic medium (pH 5.6)** |
|---|---|---|---|
| **Zero Order** | $R^2$ | 0.85 | 0.90 |
| | AIC | 114.34 | 125.56 |
| | $k$ | 0.60 | 1.14 |
| **First Order** | $R^2$ | 0.88 | 0.94 |
| | AIC | 112.23 | 105.10 |
| | $k$ | 0.23 | 0.45 |
| **Highuchi** | $R^2$ | 0.97 | 0.98 |
| | AIC | 95.12 | 89.04 |
| | $k$ | 4.98 | 9.18 |
| **Hixson-Crowell** | $R^2$ | 0.87 | 0.95 |
| | AIC | 105.45 | 98.56 |
| | $k$ | 0.64 | 1.14 |
| **Korsmeyer-Peppas** | $R^2$ | **0.99** | **0.98** |
| | AIC | 73.32 | 68.54 |
| | $k$ | 9.28 | 6.64 |
| | $n$ | 0.37 | 0.50 |
| **Peppas-Sahlin** | $R^2$ | **0.99** | **0.99** |
| | AIC | 70.34 | 63.41 |
| | $k_1$ | 10.09 | 11.01 |
| | $k_2$ | -0.005 | -0.011 |
| | $m$ | 0.43 | 0.85 |

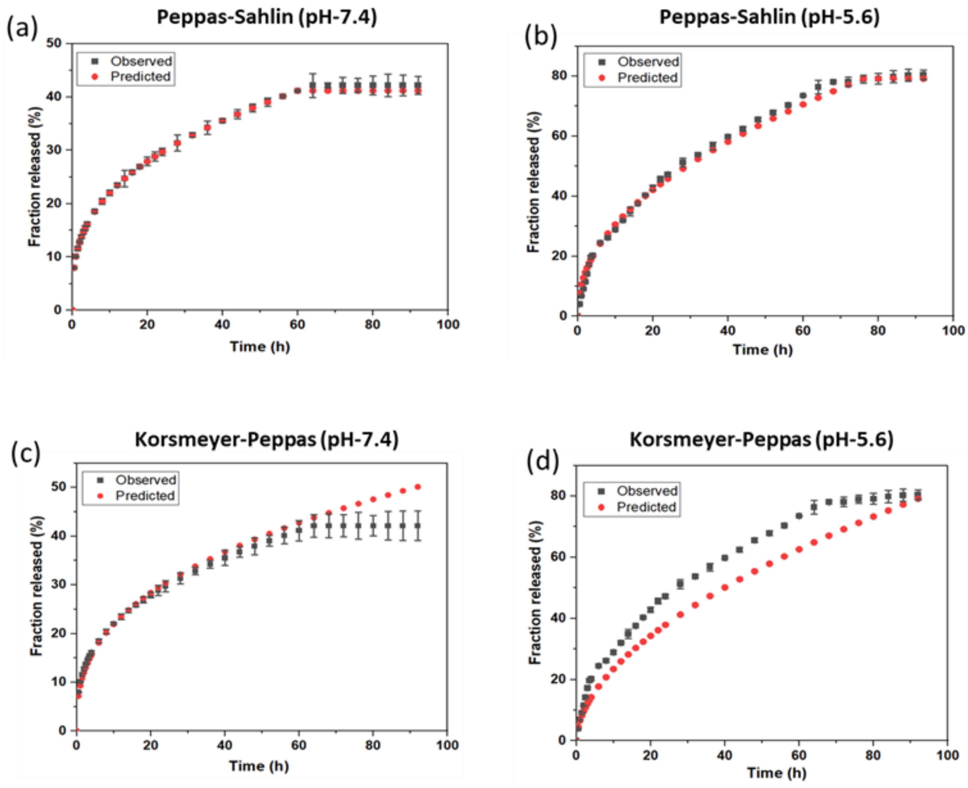

**Figure S4.** Model fitting analysis based on the Cip release profiles from Cur-EPS micelle at the two different release microenvironments (pH 7.4 at 37 °C; pH 5.6 at 37 °C) (a and b) Peppas-Sahlin, (c and d), and Korsmeyer-Peppas respectively.

**Appendix S6:**

**Storage stability of Cip-Cur-EPS**

The storage stability of Cip-Cur-EPS in PBS at 4 °C are presented in Fig. S5. The hydrodynamic diameter, PDI and EE % of Cip-Cur-EPS micelles remained consistent over the storage period for two weeks. These observations suggests that the Cip loaded micelles possesses commendable storage stability at 4°C.

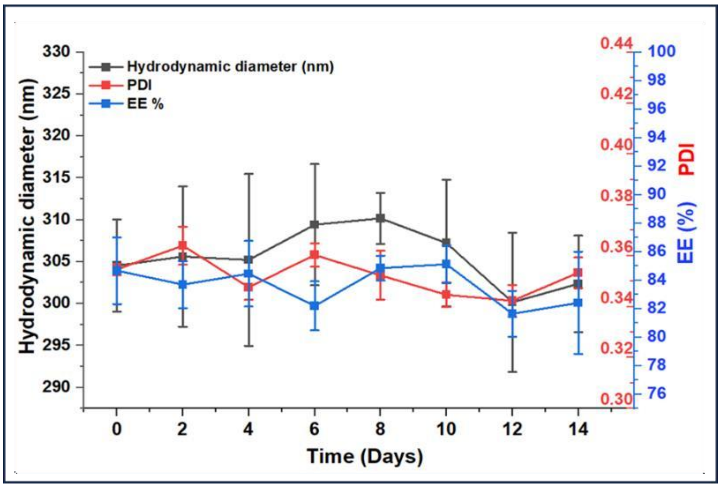

**Figure S5.** The storage stability of Cip-Cur-EPS in terms of hydrodynamic diameter, EE % and PDI until up to 14 days at 4 °C. The values are represented as mean ± SD, n = 3.

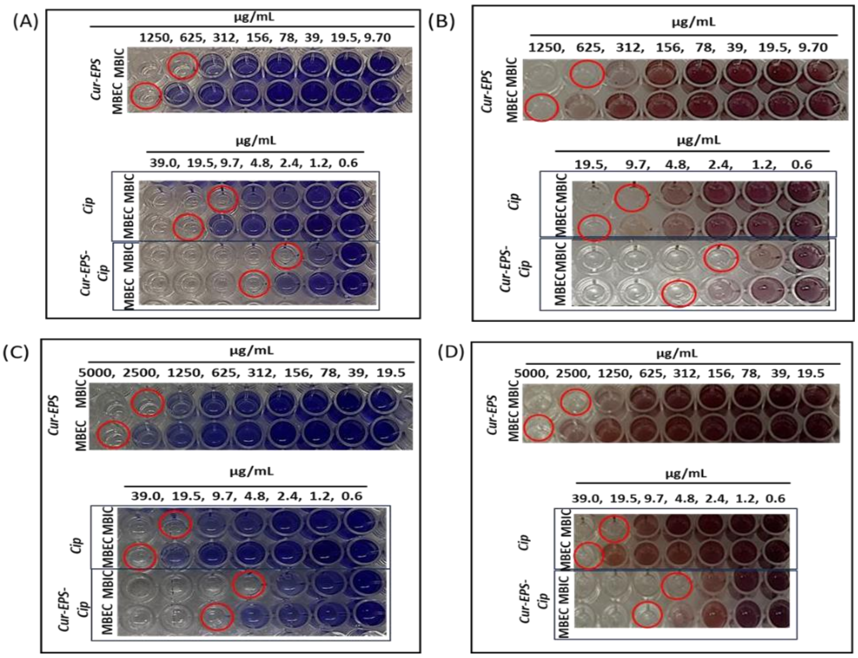

**Figure S6.** CV assay to determine concentration dependent inhibition and eradication of biofilm biomass formed by (A) *P. aeruginosa* and (C) MRSA. TTC assay to determine concentration dependent inhibition and eradication of viable bacteria within the biofilm formed by (C) *P. aeruginosa* and (D) MRSA.

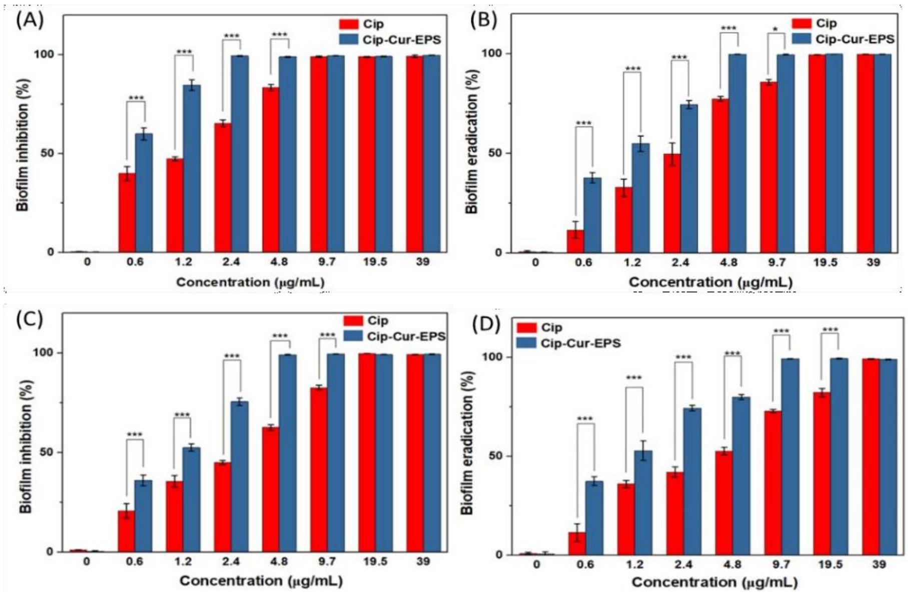

**Figure S7.** Biofilm inhibition percentage of blank Cur-EPS, free Cip, and Cip-Cur-EPS against *P. aeruginosa* (A) and MRSA (C). Biofilm eradication percentage of blank Cur-EPS, free Cip, and Cip-Cur-EPS against *P. aeruginosa* (B) and MRSA (D). Statistically significant difference ($p < 0.001$ are denoted by ***), values are represented as mean ± SD, n = 3.

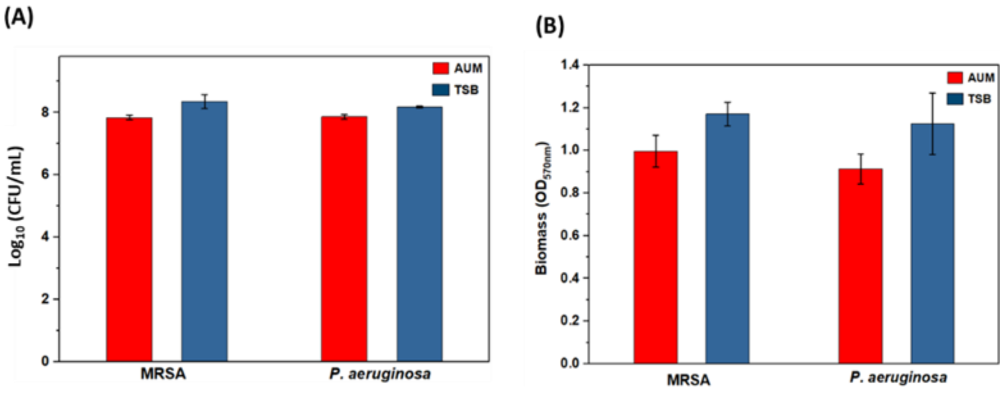

**Figure S8.** (A) Growth of MRSA and *P. aeruginosa* in AUM and TSB after 24 h at 37 °C. (B) Biofilm biomass formation by MRSA and *P. aeruginosa* in AUM and TSB after 72 h at 37 °C. The values are represented as mean ± SD, n = 3.
